# Pigs lacking Natural Killer T cells have altered cellular responses to influenza

**DOI:** 10.1101/2025.10.23.684238

**Authors:** Taeyong Kwon, Weihong Gu, Igor Morozov, Mariano Carossino, Eu Lim Lyoo, Chester D. McDowell, Yonghai Li, Udeni B.R. Balasuriya, Yi Huang, Kiho Lee, Juergen A. Richt, John P. Driver

**Affiliations:** Department of Diagnostic Medicine/Pathobiology, College of Veterinary Medicine, Kansas State University, Manhattan, KS 66506, USA; Division of Animal Sciences, University of Missouri, Columbia, MO 65211, USA; Bond Life Sciences Center, University of Missouri, Columbia, MO 65211, USA; Louisiana Animal Disease Diagnostic Laboratory and Department of Pathobiological Sciences, School of Veterinary Medicine, Louisiana State University, Baton Rouge, LA 70803, USA; National Swine Resource and Research Center, University of Missouri, Columbia, MO 65201, USA

**Author notes:** Correspondence (J.A.R.); (J.P.D.).

## Abstract

It is increasingly recognized that innate T cells such as natural killer T (NKT) cells, mucosal associated invariant T (MAIT) cells, and γδ T cells play an important role in shaping adaptive immune responses following influenza infection or vaccination. This is largely through the multiple cytokines these cells release upon activation, which have downstream effects on the scope and magnitude of virus-specific T and B cells, and antibodies which form. Here, we examined the contribution of NKT cells using pigs, which are considered a highly translational model of human influenza A infection. *CD1D*-expressing and *CD1D*-deficient pigs that respectively possess and lack NKT cells were infected with the swine influenza virus H3N2 A/Swine/Colorado/23619/1999 (CO99), with or without prior mucosal immunization with a recombinant H3N2 A/Swine/Texas/4199-2/1998 (TX98) modified live vaccine encoding a truncated NS1 protein (TX98 NS1Δ126). Vaccination reduced virus load and pulmonary pathology by similar amounts in both genotypes. However, NKT cells status had a significant impact on the underlying immune response: Unlike following vaccination, virus-specific T cell expansion after infection was greater in *CD1D*-deficient than -intact pigs, indicating that NKT cells play opposing roles in different phases of the immune response. NKT cell-deficient pigs also had reduced T cells cuffing around airways and higher numbers of expanded T and B cell clones according to single-cell αβ T cell receptor (TCR) and B cell receptor (BCR) sequencing. Using our newly developed porcine-specific γ and δ chain primers, we characterized the pulmonary γδ TCR repertoire. This revealed a high frequency of expanded clones, especially within the CD2^−^ subset, and a tendency for larger expansion in NKT cell-expressing pigs. Overall, our results indicate a homeostatic role for NKT cells on several important features of the influenza immune response, including the dynamics of T cell expansion and contraction.

## Introduction

Invariant natural killer T (NKT) cells are a population of innate T cells expressing a semi-invariant T cell receptor (TCR) that recognizes lipid and glycolipid ligands presented by the major histocompatibility complex (MHC) class I-like molecule CD1d (1–3). Upon activation, NKT cells supply a wide array of helper functions analogous to CD4^+^ T helper cells, which can enhance cellular and humoral immune responses against a range of pathogens, including against influenza A virus (IAV) (4). Among the downstream responses modulated by activated NKT cells is the licensing of antigen presenting cells that subsequently shape CD4^+^ and CD8^+^ T cell effector and memory functions (5). NKT cells also boost humoral immunity by either directly interacting with B cells presenting glycolipids via CD1d (known as cognate help), or by indirectly activating B cells via inducing T follicular helper (Tfh) cells specific for the B cell-displayed protein antigen (known as non-cognate help) (6–9).

Synthetic NKT cell ligands, such as α-GalCer, have been useful in understanding NKT cell-mediated immune responses as they specifically activate NKT cells to induce potent immunity against a wide range of co-delivered antigens. Much of this research is based on immunizing mice with IAV vaccines, with and without α-GalCer. These studies almost invariably find that NKT cell stimulation leads to stronger vaccine-induced immunity and greater protection against live virus challenge with both homologous and heterologous strains (3). However, the question of whether unmanipulated NKT cells have a natural role to play in influenza immunity is less certain. Prior studies reported that NKT cell-deficient *CD1d*- and *J*α*18*-knockout (KO) mice produced lower concentrations of influenza-specific antibodies and/or T cells following influenza infection or vaccination compared to standard mice (10, 11). In contrast, NKT cells have been reported to play a key role in suppressing influenza-specific CD8^+^ T cells through indoleamine 2,3-dioxygenase (IDO), an important mediator of immune suppression (12). In another study, *CD1d*-KO mice previously infected with H1N1 or H3N2 subtypes of IAV and re-infected after four weeks with homologous or heterosubtypic viruses were found to be just as resistant to re-infection as standard mice, indicating that, at a practical level, NKT cell responses were superfluous (13).

While these and other mouse studies have provided important insight into the role of NKT cell in influenza immunity, caution is needed in extrapolating their findings to humans, or other species with the CD1d-NKT cell system, as there are considerable inter-species differences in NKT cell frequencies and subsets (14–23). Moreover, mice are not naturally infected by IAVs and usually develop worse clinical disease than humans, but without IAV-specific clinical signs (24–27). Accordingly, it would be of benefit to re-examine the natural role of NKT cells in IAV-specific immunity using species like pigs, which are both natural IAV hosts and that share many anatomical, physiological, and immunological traits with humans, including similar NKT cell frequencies.

Hence, the goal of the current study was to compare pre-existing IAV immunity in *CD1D*-deficient and *CD1D*-expressing pigs that respectively lack and possess NKT cells. The pigs were immunized with a modified live IAV vaccine and subsequently challenged with a heterologous IAV to test cross-protective immunity. The results were compared to infected naive pigs of the same genotypes, which is of interest because NKT cells have been shown to participate in early innate immune responses that inhibit virus replication in mice (28–31). Our results shed light on how NKT cells coordinate the immune response to IAV in an animal model that closely mirrors human IAV infections.

## Materials and methods

### Virus and vaccine preparation

The modified live virus (MLV) vaccine was generated by reverse genetics from H3N2 A/Swine/Texas/4199-2/1998 (TX98) influenza virus as described previously (32). Briefly, the MLV (TX98 NS1Δ126) encodes a truncated NS1 protein with four stop codons introduced after 126 reading codons, resulting in a 3’ truncation of the wild-type NS1 protein from 219 to 126 amino acids. The challenge virus, H3N2 A/Swine/Colorado/23619/1999 (CO99), has previously been described (33). Both the vaccine and challenge viruses were propagated on MDCK cells in IAV infection media (DMEM supplemented with 0.3% bovine serum albumin, 1% MEM vitamin, 1% antibiotic-antimycotic solution and 1µg/mL of TPCK-treated trypsin).

### Pigs

The National Swine Resource and Research Center (NSRRC) at University of Missouri produced piglets that were homozygous (*CD1D−/−*) and heterozygous (*CD1D−/+)* for a 1,598 bp deletion in the *CD1D* gene, which has been described (34, 35). This *CD1D* breeding herd is on a commercial Large White crossbred background and maintained under specific pathogen free conditions. The *CD1D* genotypes of pigs were determined by PCR as previously described (Figure S1) (34, 35). The studies were in accordance with Kansas State University’s Institutional Animal Care and Use Committee (project number 4708) and Institutional Biosafety Committee (protocol number 1757).

### Experimental design

Eleven *CD1D−/−* and 14 *CD1D−/+* pigs were transferred to biocontainment rooms at 4 weeks of age after being confirmed seronegative for H1 and H3 antibodies by a hemagglutination inhibition assay, as previously described. After a 5-day acclimatization period, the pigs were assigned to 1 of 5 treatment groups. On day 0, 6 *CD1D−/−* and 6 *CD1D−/+* pigs in group (G)1 (G1) and G2, respectively, were intranasally vaccinated with 10^6^ median (50%) of tissue culture infectious dose (TCID_50_) of TX98 NS1Δ126 using an atomization device (MAD Nasal™, Teleflex, Morrisville, NC, USA). At the same time, 5 *CD1D−/−* and 5 *CD1D−/+* pigs in G3 and G4, respectively, were left unvaccinated. G5, which served as a control, contained 3 *CD1D−/+* pigs that were left unvaccinated and subjected to postmortem examination at 17 days post vaccination (DPV) after sedation with tiletamine–zolazepam (Telazol®; 4.4 mg/kg of body weight) and xylazine (2.2 mg/kg) and euthanasia with pentobarbital sodium IV injections (100 mg/kg of body weight). At 21 DPV [0 days post-challenge (0 DPC)], G1–4 were intratracheally challenged with 10^6^ TCID_50_ of CO99 as previously described and monitored for 5 days (33). Peripheral blood was collected from the jugular vein into heparin-coated or serum collection vacutainer tubes (BD Biosciences, San Jose, CA, USA) at -1, 14, 20, and 26 DPV. In order to isolate white blood cells (WBC), peripheral blood was treated with an ammonium chloride-based lysis buffer to remove red blood cells (RBC) (36, 37). Peripheral blood mononuclear cells (PBMCs) were isolated from blood samples by density gradient centrifugation using Ficoll-Paque™ PREMIUM (GE Healthcare BioSciences Corp., Uppsala, Sweden) and SepMate™ tubes (STEMCELL Technologies, Cambridge, MA, USA). Cells were cryopreserved in liquid nitrogen until use. Nasal swabs were collected at -1, 1, 3, and 5 DPV from vaccinated groups (G1 and G2) and 20, and 22–26 DPV (-1 and 1–5 DPC) from infected pigs (G1–4) in 2 mL DMEM (Corning, Corning, NY) supplemented with 1× antibiotic-antimycotic (Gibco Life Technologies, Carlsbad, CA), filtered using a 0.45 μm syringe filter (TPP, Trasadingen, Switzerland) and stored at −80 °C. At 5 DPC (26 DPV), infected pigs (G1–4) were euthanized as described above. During necropsy, bronchioalveolar lavage fluid (BALF) was collected in 50mL of DMEM supplemented with antibiotic-antimycotic (Gibco Life Technologies, Carlsbad, CA), and tissue samples, including nasal turbinates, trachea, lung, and tracheobronchial lymph nodes (TBLNs), were collected for virological, flow cytometric, and pathological evaluation. Lung cells were isolated from approximately 3 g of cranial, middle and caudal lung lobes (1 g per each lobe) as previously described with minor modifications (33): lung tissues were enzymatically digested with 2.5 mg/mL Liberase TL (Roche), 10 mg/mL DNase (Sigma), and 50 mg/mL collagenase (Worthington) in DMEM at 37°C for 30 min and mechanically dissociated using a gentleMACS™ Octo Dissociator (Miltenyi Biotec). Single cells from TBLN were isolated by mechanical dissociation using a gentleMACS™ Octo Dissociator and passed through a 100 µm cell strainer (Miltenyi Biotec). Cells were immediately cryopreserved in freezing media in temperature-controlled freezing containers at 1 to 2 × 10^7^ cells per/mL and stored in liquid nitrogen until used for flow cytometry, ELISpot, or single cell transcriptomics.

### Virus titration

Nasal turbinates, trachea, and lung tissue were mechanically homogenized in DMEM using a TissueLyser II (Qiagen, Germantown, MD) and stainless-steel beads. The resulting 10% (w/v) tissue homogenates were filtered through a 0.45 μm syringe filter (TPP, Trasadingen, Switzerland). A 10-fold serial dilution of nasal swab, BALF and tissue homogenate were prepared in influenza infection media and transferred on pre-washed, confluent MDCK cells in a 96-well plate. On day 2, the plate was fixed, incubated with mouse anti-NP antibodies (HB65 hybridoma supernatant; ATCC, Manassas, VA, USA), and subsequently incubated with goat anti-mouse IgG antibodies conjugated with Alexa Flour 488 (Invitrogen, Carlsbad, CA, USA). The virus-infected cells were visualized on an EVOS FL microscope. Viral titers were determined by the Reed-Muench method (38) and expressed as log transformed values of TCID_50_/mL or TCID_50_/g, as appropriate, according to our prior publications (33, 36).

### Antibody detection and quantification

Influenza-specific antibodies in serum and BALF were determined by hemagglutinin inhibition (HI) and ELISA assays. The HI assay was performed as previously described (36). Briefly, serum was treated with receptor-destroying enzyme II according to the manufacturer’s instructions. Then, samples were serially diluted two-fold, starting 1:10 dilution, and incubated with wild-type TX98 and CO99, followed by incubation with 0.5% washed chicken RBCs as previously described (33, 36). The highest sample dilution that inhibited virus-induced RBC hemagglutination is presented.

Virus-specific IgG, IgG1, IgG2a, and IgA antibody responses were measured in serum and BALF using an in-house ELISA. Briefly, 96-well ELISA plates were coated overnight with 2 µg/mL of H3 protein from A/California/07/2004 H3N2 (Sino biologicals, Beijing, China) dissolved in coating buffer. The amino acid sequence identities of this antigen to the HA proteins of TX98 and CO99 are 91.5% and 94.9%, respectively. H3 antigen solution was removed, and plates were incubated with blocking buffer (3% goat serum, 0.5% skim milk, and 0.1% Tween 20 in PBS) at room temperature for 1 h. Plates were washed and incubated for 2 h with serum or BALF serially diluted two-fold in blocking buffer, starting from 1:100 dilution for serum and 1:10 dilution for BALF. Before preparing serial dilutions, BALF was incubated with the equal amount of 10 mM DTT for 1 h at 37°C for mucus disruption. To determine high-affinity IgG titer in serum, low-affinity antibodies were removed after incubation of 6 M urea for 10 minutes at room temperature. Plates were washed and incubated for 1 h at room temperature with 100 µL of blocking buffer containing the following isotype-specific secondary antibodies: HRP-conjugated anti-pig IgG (Invitrogen, Carlsbad, CA, USA; 1:40,000 dilution for both serum and BALF), HRP-conjugated anti-pig IgA (Invitrogen, Carlsbad, CA, USA; 1:20,000 dilution for serum and 1:40,000 for BALF), anti-pig IgG1 (Bio-Rad, Hercules, CA, USA; 1:1000 dilution), and anti-pig IgG2 (Bio-Rad, Hercules, CA, USA; 1:5,000 dilution). For the IgG1 and IgG2 ELISAs, plates were further incubated with an HRP-conjugated goat anti-mouse IgG (H+L) antibody (Invitrogen, Carlsbad, CA, USA: 1:20,000 dilution) for 1 hour. Plates were washed and incubated with 3,3’,5,5’ tetramethylbenzidine solution (Thermo Scientific™,

Rockford, IL, USA) for 15 min before adding stop solution (Abcam, Cambridge, MA, USA) and read at 450nm using an ELISA plate reader. The detection cut-off was calculated as the average optical density (O.D.) + 3 × standard deviations of the three G5 control pigs. ELISA titers are represented as the reciprocal of the highest serum dilution above the cut-off value.

### Flow cytometry

After thawing, approximately 1-2 million WBC, TBLN, and lung cells were stained using our previously described protocols, with minor modifications (33). Briefly, cell suspensions were incubated with a viability dye (LIVE/DEAD™ Fixable Near-IR Dead Cell Stain Kit, Invitrogen, Carlsbad, CA, USA), followed with Fc blocking with rat IgG (Sigma-Aldrich, St. Louis, MO, USA), and then a panel of monoclonal antibodies to identify immune cell populations. T cell and NK cell subsets were identified using antibodies specific for CD3ε (BB23-8E6-8C8; BD Biosciences, Franklin Lakes, NJ, USA), CD4 (74-12-4; Southern Biotech, Birmingham, AL, USA), CD8α (76-2-11; Southern Biotech), CD8β (PPT23; Bio-Rad, Hercules, CA, USA), TCRδ (PGBL22A; WSU Monoclonal Antibody Center, Pullman WA, USA), CD16 (G7; BD Biosciences, Franklin Lakes, NJ, USA), and CD11b (M1/70; BioLegend, San Diego, CA, USA). NKT cells were identified using unloaded and PBS57-loaded mouse CD1d tetramers provided by the National Institutes of Health Tetramer Core Facility. Monocytes, macrophages, and granulocytes were distinguished using antibodies against CD14 (MIL2; Bio-Rad, Hercules, CA, USA), CD16, CD163 (2A10/11; Bio-Rad, Hercules, CA, USA), CD172a (74-22-15A; BD Biosciences, Franklin Lakes, NJ, USA), CD11b, and MHC class II (H42A; WSU Monoclonal Antibody Center). After staining, cells were fixed using the BD Cytofix/Cytoperm kit (BD Biosciences) and acquired on a BD LSRFortessa™ X-20 flow cytometer with FACSDiva software (version 9.2, BD Biosciences).

Fluorescence-minus-one controls were used to determine positive and negative populations. Data was analyzed using FlowJo software (version 10.10.0, Treestar, Palo Alto, CA, USA). Leukocyte populations were identified using the gating strategy as previously published (33).

### ELISpot Assays

After thawing, PBMCs and lung cells were resuspended in RPMI supplemented with 10% FBS, 1% antibiotic-antimycotic, and 55 μM 2-mercaptoethanol (Gibco, Waltham, MA, USA) and rested for at least 1 h. For the IFN-γ ELISpot assay, PBMCs and lung cells were respectively plated at 2.5×10^5^ and 2×10^5^ cells per well in 96-well MultiScreen IP HTS plates (Millipore, Billerica, MA, USA) pre-coated with anti-IFN-γ (P2G10, BD Biosciences, Franklin Lakes, NJ, USA). The cells were incubated for 48 h with 0.1 MOI of wild-type TX98 and CO99 virus or virus-free MDCK supernatant. Plates were then developed with a biotin-conjugated anti-IFN-γ mAb (P2C11, BD Biosciences, Franklin Lakes, NJ, USA), streptavidin-horseradish peroxidase (BD Biosciences, Franklin Lakes, NJ, USA), and 3-amino-9- ethylcarbazole (AEC) substrate (BD Biosciences, Franklin Lakes, NJ, USA), according to manufacturer instructions. A similar HRP-based IL-2 ELISpot assay was performed using 2.5×10^5^ PBMCs per well stimulated with 0.1 MOI of wild-type TX98 and CO99 virus or virus-free MDCK supernatant, as per the manufacturer’s instruction (Mabtech, Stockholm, Sweden). The number of spots in each well were counted using an AID iSpot EliSpot FluoroSpot Reader with AID EliSpot Software Version 7.0 (Advanced Imaging Devices GmbH, Strassberg, Germany).

### Gross pathology, histopathology, and immunohistochemistry

At necropsy, the apical and cardiac lung lobes were fixed in 10% neutral buffered formalin, processed and embedded in paraffin, and stained with hematoxylin and eosin (H&E) for histological evaluation following standard procedures. At least two sections of lung per animal were blindly scored for histological lesions according to our previous publication, with minor modifications (39): (i) epithelial necrosis, attenuation, disruption or hyperplasia (0–4), (ii) airway exudate (0–4), (iii) percentage of airways with inflammation (0–4), (iv) peribronchiolar and perivascular lymphocytic inflammation (0–3), (v) alveolar exudate (0–3), and (vi) alveolar septal inflammation (0–4).

For abundance and distribution of CD3^+^ T cells in the pulmonary parenchyma, 4-micron formalin-fixed, paraffin-embedded (FFPE) tissue sections were subjected to immunohistochemistry for CD3 using the automated BOND RXm platform and the Polymer Refine Detection kit (Leica Biosystems, Buffalo Grove, IL). Following automated deparaffinization, FFPE tissue sections were subjected to automated heat-induced epitope retrieval (HIER) using a ready-to-use citrate-based retrieval solution (pH 6.0, Leica Biosystems) at 100 °C for 20 min. Subsequently, tissue sections were incubated with the primary antibody (rabbit polyclonal anti-human CD3 [A0452, Dako, Carpinteria, CA] diluted 1:300 in Primary Antibody diluent [Leica Biosystems]) for 30 min at ambient temperature, followed by a polymer-labeled goat anti-rabbit IgG coupled with HRP (8 min). 3’,3’ diaminobenzidine (DAB) was used as the chromogen (10 min), and counterstaining was performed with hematoxylin for 5 min. Slides were dried in a 60°C oven for 30 min and mounted with a permanent mounting medium (Micromount®, Leica Biosystems). A pathologist blinded to the study design evaluated abundance of CD3^+^ T cells (0, none; 1, minimal numbers; 2, mild numbers; 3, moderate numbers; 4, high numbers). In addition, CD3^+^ distribution in the pulmonary parenchyma of each pig was classified into two categories: (1) scattered in pulmonary parenchyma with no defined cuffing around airways or (2) airway-centric distribution cuffing small and larger airways.

### Single-cell processing

Thawed lung cells were used to generate libraries as previously described (40). Gene expression libraries were prepared using the Chromium Next GEM Single Cell 5′ Kit v2 (10× Genomics, Pleasanton, CA) according to the manufacturer’s instructions. V(D)J libraries were enriched for αβ TCR and B cell receptor (BCR) transcripts with pig-specific primers, as previously described (40). We also included new porcine-specific γ and δ chain TCR primers (Table S1), which were designed and run according to our previously developed single-cell αβ TCR and BCR assays (40).

### Single-cell RNAseq (scRNAseq) analysis

The Sscrofa 11.1 genome assembly was used to align sequencing reads to generate gene matrix data by Cell Ranger (v8.0.0). Each dataset was pre-processed by removing genes expressed in <3 cells, excluding cells with aberrantly high or low gene counts and high mitochondrial gene expression. Afterwards, batch correction and clustering analyses were performed using Seurat (v.5.3.0) (41). Briefly, transcript counts were log normalized, and highly variable genes were selected for dimensionality reduction analysis. The IntegrateLayers function was used to align cells across multiple samples by correcting batch effects while preserving biological variability, using the CCA integration method. Then, the clustering analysis workflow was performed using FindNeighbours, FindClusters, and RunUMAP functions. The FindTransferAnchors and TransferData functions were used to infer the cell types from our previous datasets (40), and further validated based on the expression of known cell type markers. The differentially expressed genes (DEGs) between treatments in each cluster were computed using FindMarkers function with the Wilcoxon test. The pathway enrichment analysis was performed using Ingenuity Pathway Analysis (IPA) (QIAGEN Inc.). The Core Analysis function was used to identify canonical pathways significantly associated with the DEGs using a p-value threshold of 0.05.

### Single-cell V(D)J data analysis

Single-cell αβ and γδ TCR and BCR V(D)J sequencing reads were assembled into contigs using the cellranger vdj (10× Genomics) pipeline in de novo mode. To identify the V(D)J chains, we searched assembled contigs against inner-enrichment primers using the usearch_global command. The TCR β, γ, and BCR V(D)J chains were respectively mapped to the pig TRB and IG reference sequences in IMGT using the IMGT/V-QUEST (42). Using MMseqs2 (v17.b804f) (43), primer-matched TCR α TRAJ and TCR δ segments were further mapped to the pig TRAJ and TRDJ IMGT genome database, respectively. TRAV and TRDV segments were annotated based on TRAV/TRDV sequences deposited in GenBank (44), which we previously named according to their similarity to human TRAV genes (45). Single cells with a single unique annotated contig in any of the TCR αβ, γδ, or BCR IGH, IGL, and IGK chains were retained for downstream analysis, while those with multiple contigs per chain were excluded. The filtered annotated contigs were then aligned to their corresponding gene expression profiles. Scirpy (v0.12.0) was used to analyze CDR3 clonal expansion, CDR3 amino acid sequence length, and V(D)J segment usage analyses across samples (46). In addition, TCR β-chain CDR3 amino acid sequences were analyzed using TCRmatch (http://tools.iedb.org/tcrmatch/) for antigen prediction. The tool compares the input β-chain CDR3 sequences against those in the Immune Epitope Database (IEDB), identifies similar sequences, and retrieves the corresponding epitopes and antigens annotated in the IEDB (47). CDR3 clonal and V(D)J segment diversity was quantified using the Alakazam package (v1.3.0) (48). Diversity was assessed across a range of diversity orders (q) based on the Hill diversity framework, with particular emphasis on the exponential Shannon-Weiner index (q=1) to evaluate V(D)J segment diversity across different cell types (49).

## Data availability

The sequencing data are available at Gene Expression Omnibus (accession GSE306556). All relevant data are available from the authors.

## Statistics

The virus titers were log-transformed for statistical analysis. Analysis of variance (ANOVA) and subsequent Tukey’s adjustment were performed for virus titers, pathological scores, HI titers, ELISPOT and flow cytometry and t-test were used for ELISA, cell frequencies from scRNA-seq analysis, as well as for clonal expansion, CDR3 length, and antibody isotypes derived from scTCR/BCR-seq analyses, using GraphPad Prism 10 (Boston, MA, USA). CD3^+^ T cell distribution was analyzed using a nominal logistic regression model in JMP Statistical Discovery (SAS, Cary, NC).

## Results

### Virus shedding and replication

Four-week-old pigs carrying an inactive form of the *CD1D* (*CD1D−/−*) gene and littermates carrying one inactivated copy (*CD1D−/+*) were intranasally vaccinated with a recombinant H3N2 A/Swine/Texas/4199-2/1998 (TX98) influenza virus encoding a truncated NS1 protein (TX98 NS1Δ126). Pigs were subsequently infected at 21 DPV with the heterologous H3N2 A/Swine/Colorado/23619/1999 (CO99) virus and monitored for 5 days. Additional groups included *CD1D−/−* and *CD1D−/+* pigs that were infected without prior vaccination, and unvaccinated *CD1D−/+* pigs that were not challenged, which were used as a negative control.

Both vaccinated groups shed similarly low levels of TX98 NS1Δ126 between 1 and 5 DPV (Figure 1A). After challenge, shedding was detected from 2 out of 6 vaccinated *CD1D−/+* pigs, while no virus was detected from the vaccinated *CD1D−/−* pigs (Figure 1B). Moreover, none of the vaccinated pigs had detectable virus in BALF, nasal turbinates, or trachea at 5 DPC (Figures 1C–E). These results indicate that the MLV vaccine was similarly effective at inhibiting virus replication in NKT cell-deficient and NKT cell-expressing pigs. Unvaccinated *CD1D−/−* and *CD1D−/+* pigs shed increasing levels of virus beginning from 1 DPC in nasal swabs and presented high virus titers in nasal turbinates, trachea, and BALF. While there was no statistical difference between the genotypes, virus titers tended to be higher in the trachea and nasal turbinates of unvaccinated *CD1D−/+* than *CD1D−/−* pigs. This suggests that, opposite to mice, NKT cells in pigs do not inhibit IAV replication and may even increase it.

**Figure 1.**
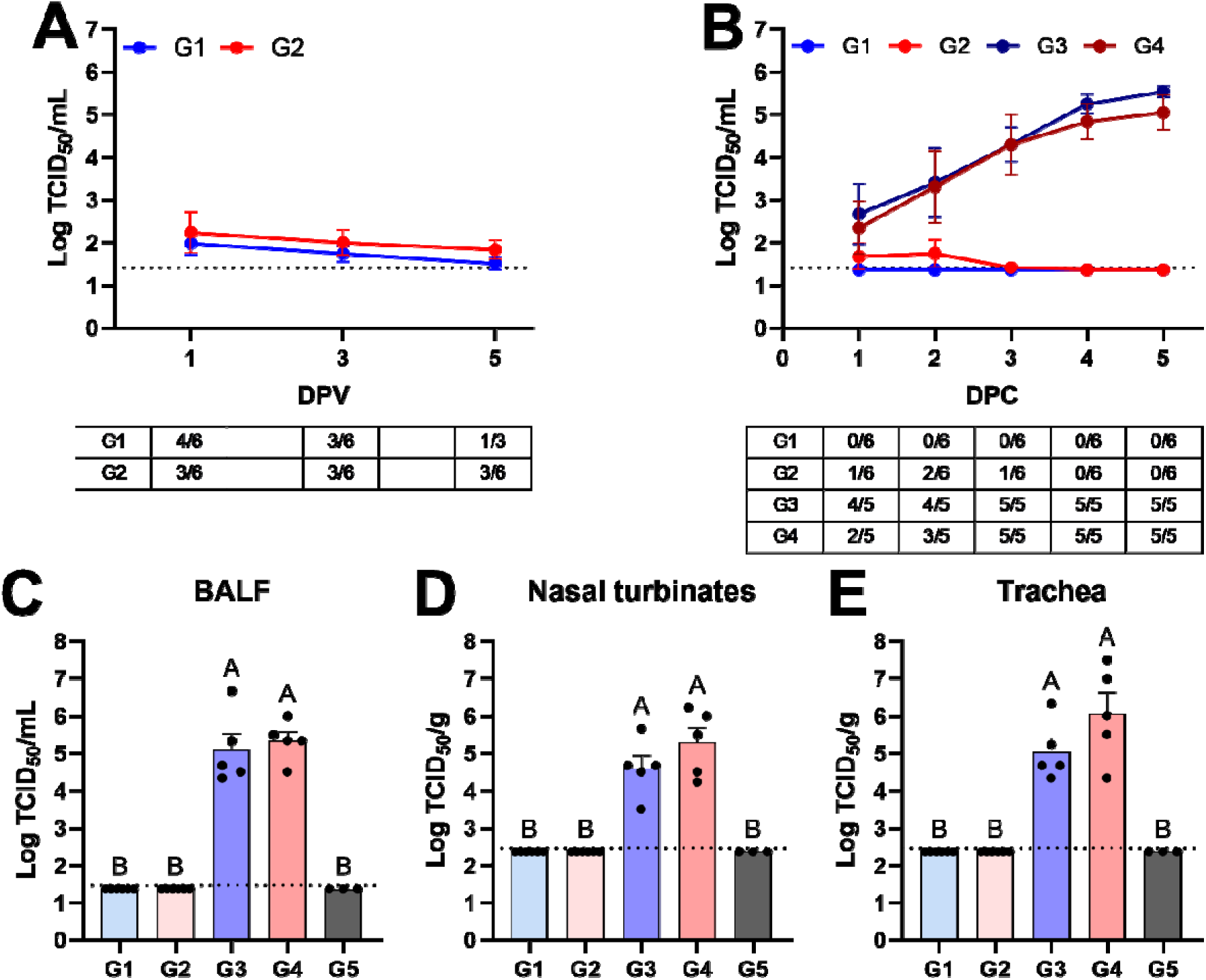
Viral titers in nasal swabs and respiratory tissues. (A, B) Virus titers in nasal swabs and frequency of pigs positive for virus shedding at 1, 3, and 5 DPV (A) and at 1–5 DPC (B). (C–E) Virus titers in bronchoalveolar lavage fluid (BALF) (C), nasal turbinates (D), and trachea (E) at 5 DPC. A statistically significant difference (*p* < 0.05) between two groups is indicated by different letters. Data are represented as mean ± SEM of log_10_ (TCID_50_/mL or TCID_50_/g). Symbols represent treatment groups (A, B) or individual pigs (C–E). G1: *CD1D−/−* vaccinated and challenged; G2: *CD1D−/+* vaccinated and challenged; G3: *CD1D−/−* not vaccinated and challenged; G4: *CD1D−/+* not vaccinated and challenged; G5: *CD1D−/+* not vaccinated, not challenged.

### Immunopathology

Microscopic changes in the pulmonary parenchyma were scored as described in the Materials and Methods section. Both vaccinated groups (G1 and G2) had overall lower cumulative scores, reflecting less pronounced IAV-associated microscopic lesions (Figures 2A and 3A–B). Unvaccinated and challenged groups (G3 and G4) had pronounced airway-centric inflammation compared to vaccinated groups (Figures 2B and 3C–D). In general, *CD1D−/−* pigs tended to have lower histopathology scores than *CD1D−/+* pigs, although this difference was not statistically significant.

**Figure 2.**
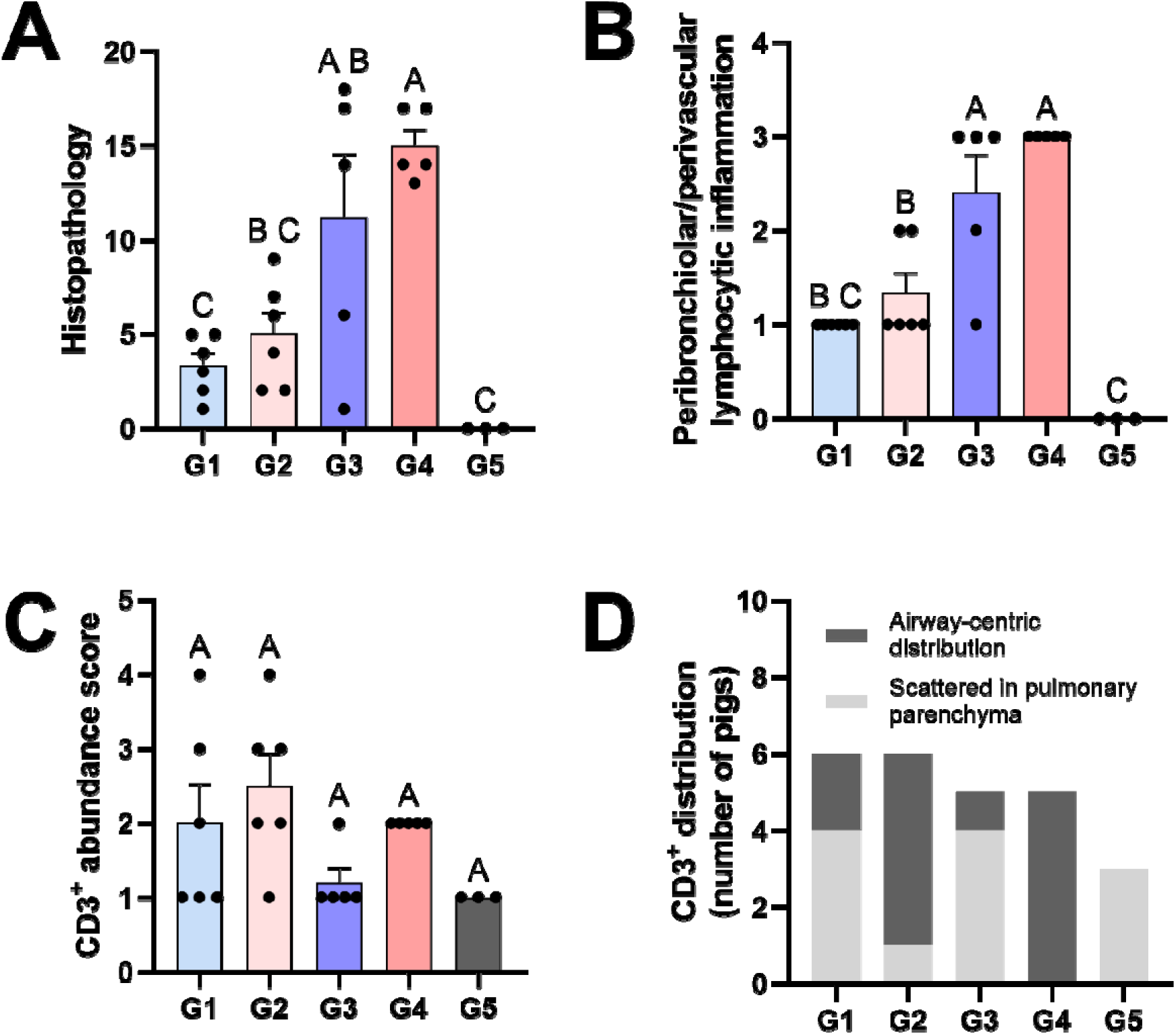
Lung histopathology and CD3^+^ T cell localization at 5 DPC. (A) Cumulative histological scores following assessment of the parameters shown in the materials and methods. Histologic alterations occurred at a similar degree between CD1D*−/+* and CD1D*−/−* pigs in the vaccinated and challenged as well as the unvaccinated and challenged groups, with the latter groups showing more severe alterations (reflected by a higher cumulative score). (B) Degree of peribronchiolar/perivascular lymphocytic inflammation. (C, D) Scores for overall CD3^+^ cell abundance in the lung (C) and distribution of CD3^+^ cells (D), assessed by immunohistochemistry. CD3^+^ distribution in the pulmonary parenchyma of each pig was classified into two categories: (1) scattered in pulmonary parenchyma with no defined cuffing around airways or (2) airway-centric distribution cuffing small and larger airways. A statistically significant difference (*p* < 0.05) between two groups is indicated by different letters (A, B, and C). Data are represented as mean ± SEM. Symbols represent individual pigs. G1: *CD1D−/−* vaccinated and challenged; G2: *CD1D−/+* vaccinated and challenged; G3: *CD1D−/−* not vaccinated and challenged; G4: *CD1D−/+* not vaccinated and challenged; G5: *CD1D−/+* not vaccinated, not challenged.

**Figure 3.**
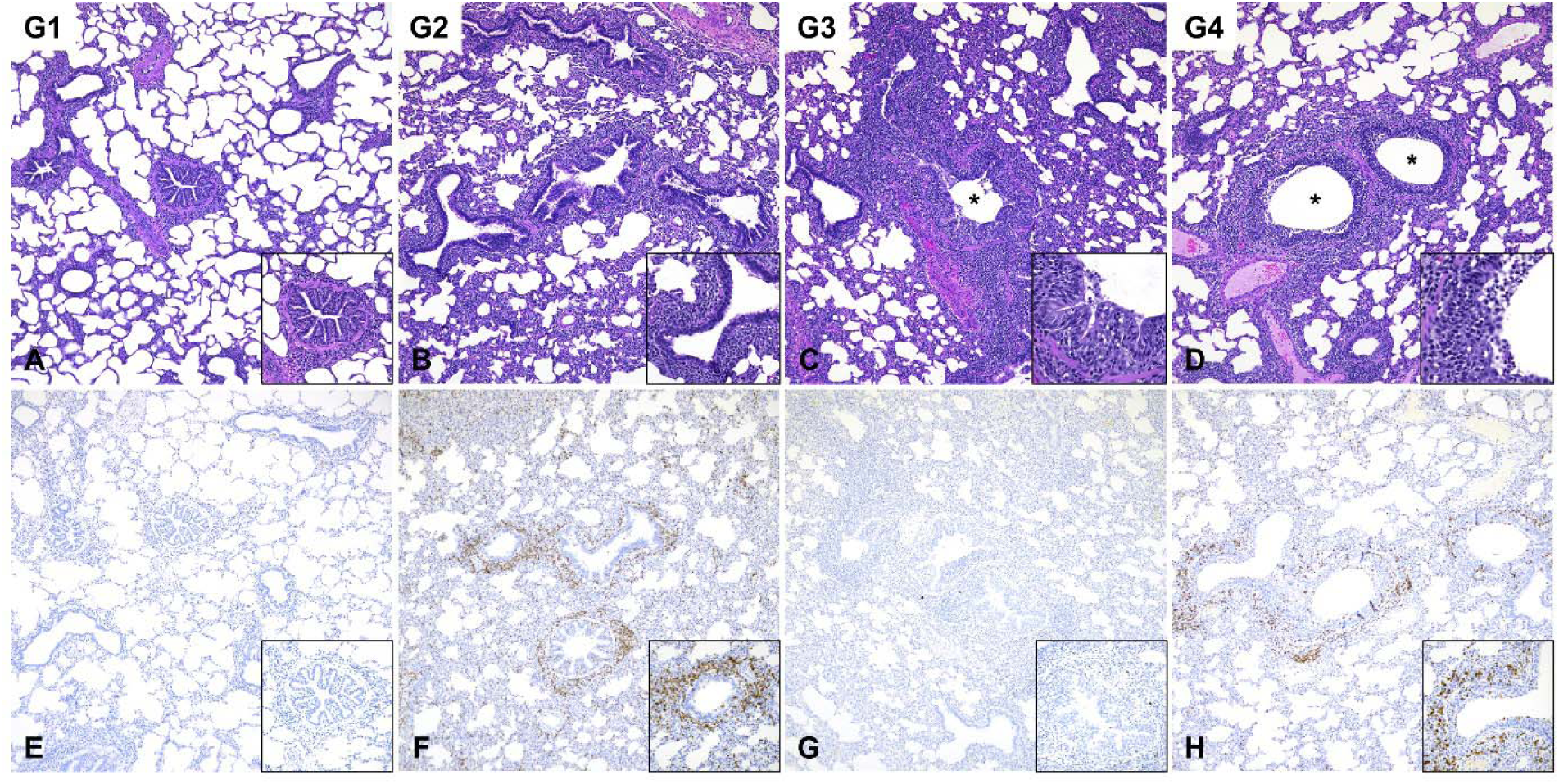
Histologic lesions (A–D) and CD3^+^ T cell distribution (E–H). Histologic alterations in G1 and G2 (A and B) are similar and featured by a mild mononuclear infiltrate delimiting bronchioles (insets) and occasionally expanding the alveolar septa. In G3 and G4 (C and D), the degree of airway-centric inflammation is higher and is additionally characterized by intraluminal exudate composed of degenerate neutrophils and a hyperplastic bronchiolar epithelium with transmigrating lymphocytes and neutrophils (asterisks and insets) and regions of pulmonary atelectasis. CD3^+^ T cells in *CD1D*−/+ pigs are intensely recruited around airways (F and H) regardless of vaccination status compared to *CD1D*−/− pigs (E and G). Immunohistochemistry for CD3 (DAB). G1: *CD1D*−/− vaccinated and challenged; G2: *CD1D*−/+ vaccinated and challenged; G3: *CD1D*−/− not vaccinated and challenged; G4: *CD1D*−/+ not vaccinated and challenged.

Subsequently, we utilized immunohistochemistry to assess whether NKT cell deficiency affected T cell localization in the lung following IAV vaccination and infection. While the overall abundance of CD3^+^ T cells was similar among the treatment groups (Figure 2C), there was a significantly greater airway-centric distribution of CD3^+^ T cells in *CD1D−/+* pigs compared to their *CD1D−/−* counterparts, regardless of vaccination status (p-value 0.0061; Figures 2D and 3E–H).

### Hemagglutinin (HA)-specific antibody responses

Vaccinated pigs had high TX98-specific HI titers throughout the vaccination and challenge periods, with no difference between genotypes (Figure 4A). While a few vaccinated pigs developed low CO99-specific HI titers during the vaccination period, all vaccinated pigs developed high CO99 titers following challenge (Figure 4B). After challenge, unvaccinated pigs developed HI titers against CO99, but not against TX98. There was no significant difference in HI titers between genotypes.

**Figure 4.**
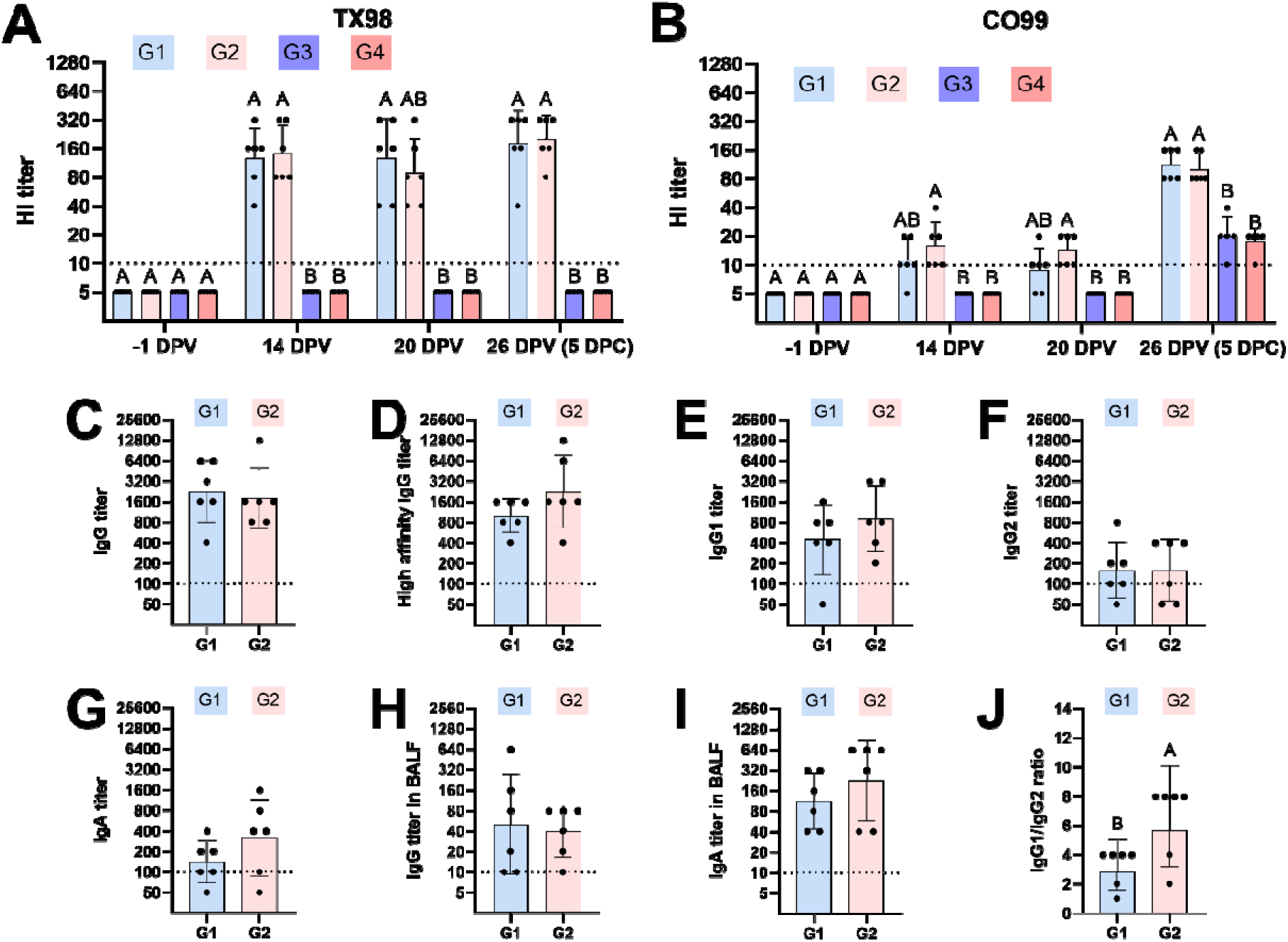
Hemagglutinin (HA)-specific antibody titers measured by hemagglutinin inhibition (HI) assay and in-house isotype-specific ELISA. (A, B) Geometric mean of HI titers against TX98 (A) and CO99 (B) in serum. (C–I) Isotype-specific (IgG, IgG1, IgG2, and IgA) antibody titers against HA in serum (C–G) and BALF (H and I) of vaccinated pigs (G1 and G2). (J) Ratio of HA-specific IgG1/IgG2 titers in serum (J). A statistically significant difference (*p* < 0.05) between two groups is indicated by different letters. Data are represented as geometric mean ± geometric SD. Symbols represent individual pigs (C-E). G1: *CD1D−/−* vaccinated and challenged; G2: *CD1D−/+* vaccinated and challenged; G3: *CD1D−/−* not vaccinated and challenged; G4: *CD1D−/+* not vaccinated and challenged; G5: *CD1D−/+* not vaccinated, not challenged.

Serum and BALF from the two vaccinated groups were measured for HA-specific IgG, high affinity IgG, IgG1, IgG2, and IgA responses at 5 DPC. Assays were performed using a HA protein that shares more than 90% amino acid identity with TX98 and CO99. End-point titers for all antibody types did not differ between genotypes (Figures 4C–I). However, *CD1D−/−* pigs had a significantly lower IgG1/IgG2 ratio when compared to *CD1D−/+* pigs (Figure 4J), suggesting that NKT cell responses affected B cell effector functions.

### Flow cytometry and single-cell RNA sequencing

Flow cytometry was used to analyze leukocyte populations in lung tissue and TBLN at 5 DPC and in blood at -1, 14, 20 and 26 DPV. Vaccinated pigs, but especially the *CD1D−/−* group (G1), had higher frequencies of CD3^+^ cells (Figure 5A) and αβ T cells (Figure 5B) as a proportion of lung lymphocytes compared to unvaccinated pigs. Furthermore, vaccination led to an increase in CD8αβ^+^ T cells as a proportion of CD3^+^ cells in lungs (Figure 5C). However, there were no notable treatment differences in other T cell subsets, natural killer (NK) cells, monocytes, macrophages, dendritic cells, or granulocytes (Table S2–S7).

**Figure 5.**
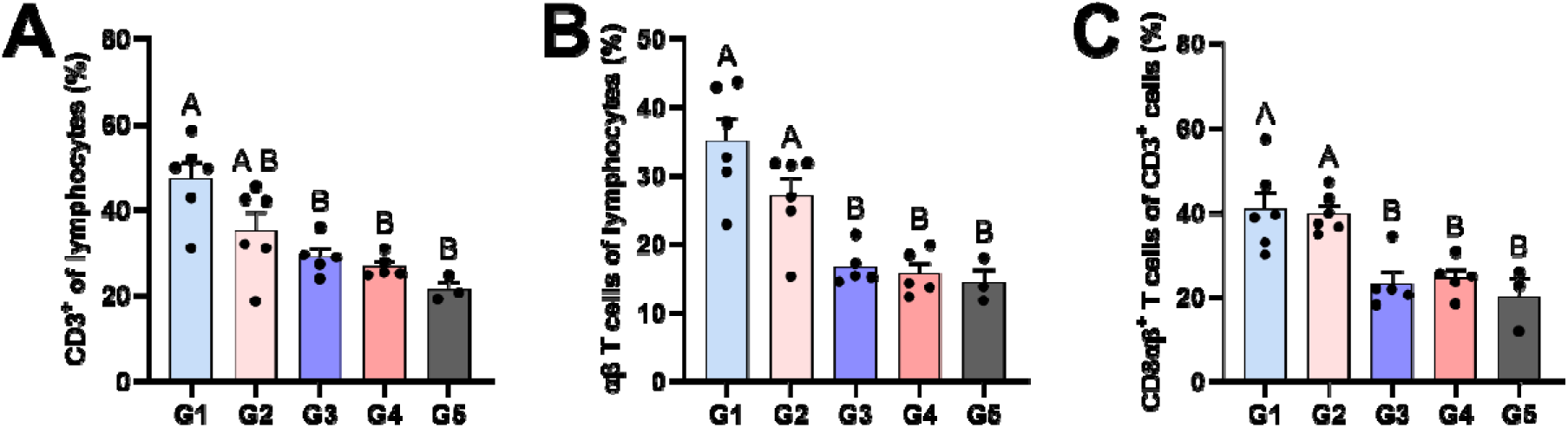
Flow cytometric analysis of T cell frequencies in enzymatically digested lung tissue at 5 DPC. (A) CD3^+^ cells as a proportion of lung lymphocytes. (B) αβ T cells as a proportion of lung lymphocytes. (C) CD8αβ^+^ T cells as a proportion of CD3^+^ cells. A statistically significant difference (*p* < 0.05) between two groups is indicated by different letters. Data are represented as mean ± SEM. Symbols represent individual pigs. G1: *CD1D−/−* vaccinated and challenged; G2: *CD1D−/+* vaccinated and challenged; G3: *CD1D−/−* not vaccinated and challenged; G4: *CD1D−/+* not vaccinated and challenged; G5: *CD1D−/+* not vaccinated, not challenged.

To obtain more detailed information on the cellular differences between genotypes, we performed single-cell transcriptomic analysis on lung tissue cells from four pigs per vaccinated group at 5 DPC, totaling 69,162 cells. A dimensionality reduction analysis identified 43 clusters by Uniform Manifold Approximation and Projection (UMAP) that we annotated according to a combination of label transfer from previous dataset (40) and established lineage markers (Figures 6A and S2A). The frequencies of CD4^+^ tissue resident memory T cells (TRMs) (cluster 3), CD2*^−^* γδ T cells (cluster 11), cycling B cells (cluster 19), and plasma cells (cluster 20) were significantly higher in *CD1D−/−* compared to *CD1D−/+* pigs (Figure 6B). The frequencies of CD8^+^ TRMs (clusters 4–6) and cycling T cells (clusters 8–10) were also higher in *CD1D−/−* pigs, but the difference was not significant. *CD1D−/+* pigs had higher numbers of resident NK cells (cluster 16) than *CD1D−/−* pigs, and a tendency for higher frequencies of cells with a mixed monocytes/macrophage phenotype (clusters 21–27) as well as monocytes (clusters 29 and 30).

**Figure 6.**
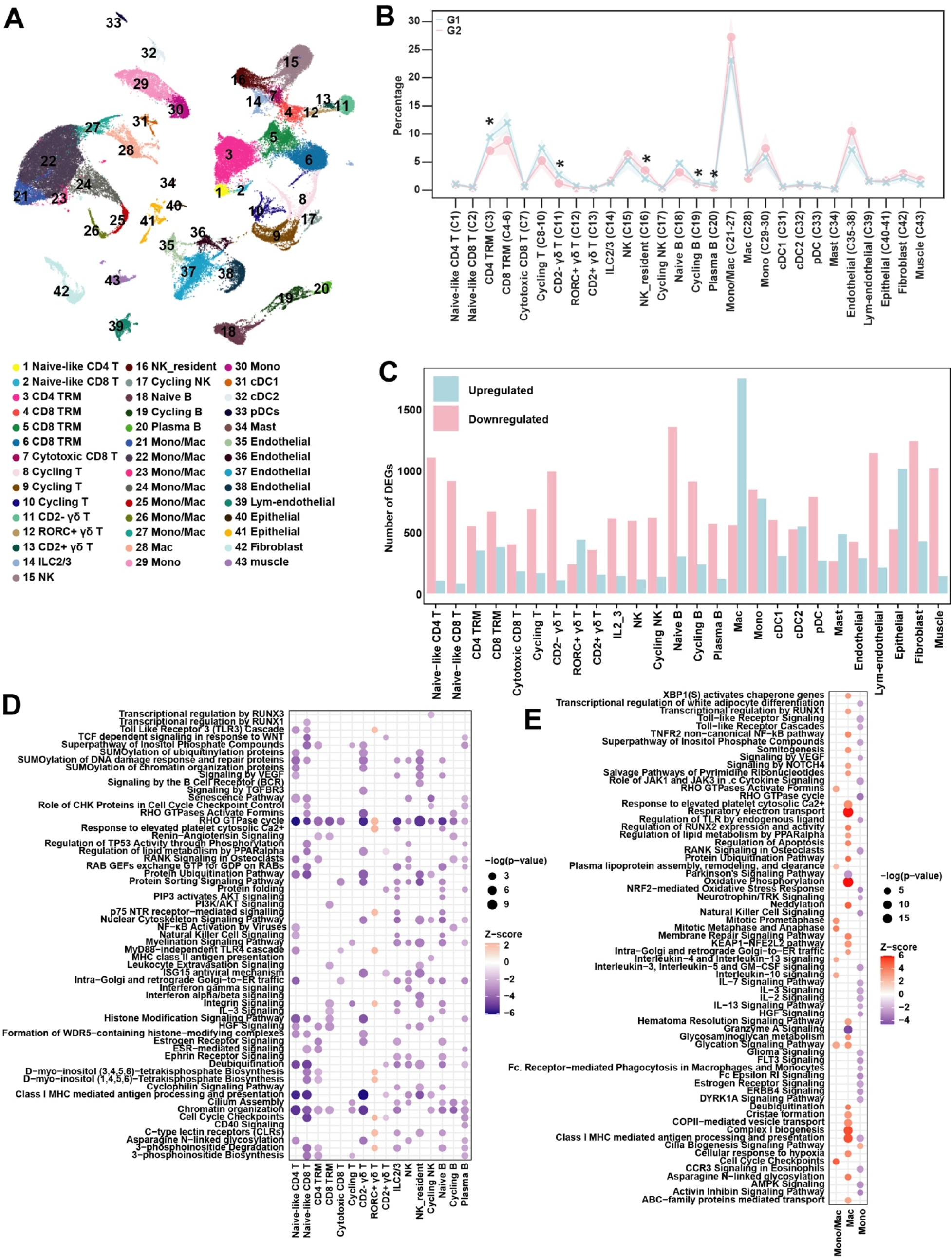
Single-cell transcriptomic analysis of vaccinated and IAV-infected *CD1D−/−* and *CD1D−/+* pig lungs. (A) UMAP visualization of pig lung cells. (B) The average frequency of each cell type is presented for each group. Significance was determined by t-test in each cell type. (C) Bar graphs displaying the number of upregulated and downregulated differentially expressed genes (DEGs) in G1 compared to G2 pigs. (D–E) Ingenuity pathway analysis (IPA) of DEGs in lymphocytes (D) and myeloid cells (E). The y-axis shows the top pathways identified in each cell type within the threshold −2.0 < Z > 2.0. Dot size indicates significance [−log_10_(P value)], and dot color saturation reflects the z-score. G1: *CD1D−/−* vaccinated and challenged; G2: *CD1D−/+* vaccinated and challenged.

Next, we compared *CD1D−/+* and *CD1D−/−* pigs for differentially expressed genes (DEGs) within individual cell types (Figure 6C, Data file 1). Overall, we detected substantially more downregulated than upregulated genes in *CD1D−/−* compared to *CD1D−/+* pigs, especially within T cell, B cell, and NK cell populations. An Ingenuity Pathway Analysis (IPA) using DEGs in each cluster revealed that *CD1D−/−* naïve CD8^+^ T cells, CD4^+^ TRMs, and plasma cells downregulated a variety of pathways involved in inositol phosphate biosynthesis and degradation (Figure 6D). Inositol phosphates play an integral role in development, proliferation, and differentiation of T and B lymphocytes (50, 51). CD8^+^ TRMs downregulated pathways related to PI3K/AKT signaling, leukocyte extravasation, integrin signaling, and IL-3 signaling. All lymphocytes, except for cycling T cells, *RORC^+^* γδ T cells, and CD2^+^ γδ T cells, downregulated the RHO GTPase cycle (Figure S2B), which is a collection of regulatory proteins critical for lymphocyte migration, polarization, adhesion, activation, and differentiation (52–55). *CD1D−/− RORC^+^* γδ T cells were the only lymphocyte population with mostly upregulated pathways compared to their *CD1D−/+* counterparts, including Toll-Like Receptor 3 (TLR3) and TLR4 cascades. *CD1D−/−* macrophages also had more upregulated than downregulated pathways than *CD1D−/+* pigs (Figure 6E). These included the class I MHC antigen processing and presentation pathway as well as oxidative phosphorylation, respiratory electron transport, and complex I biogenesis pathways, which are often enriched in macrophages undergoing a type of metabolic reprogramming associated with anti-inflammatory and healing states (56). Collectively, our findings indicate that NKT cell-derived stimuli influenced a broad range of signaling pathways and cell types in lungs of IAV-infected pigs.

### Immune receptor profiling

To study the relationship between NKT cell status and immune receptor repertoire diversity, we coupled our scRNAseq dataset with additional assays to enrich αβ TCR and BCR chains using primers that target the C regions in mRNA transcripts of αβ TCR and BCR chains and isotypes. Over the eight samples, we obtained an average of 187,771,466 reads per αβ TCR library and 206,842,207 reads per BCR library (Data file 2). Across all samples, most TRA- or TRB- positive cells were mapped to T/NK/ILC clusters (1–17), which contained an average of 95% of paired TRA+TRB cells (Figure 7A, Data file 2). Only T/NK/ILC cells with annotated paired TRA and TRB chains were used for further analysis (Figure 7B).

**Figure 7.**
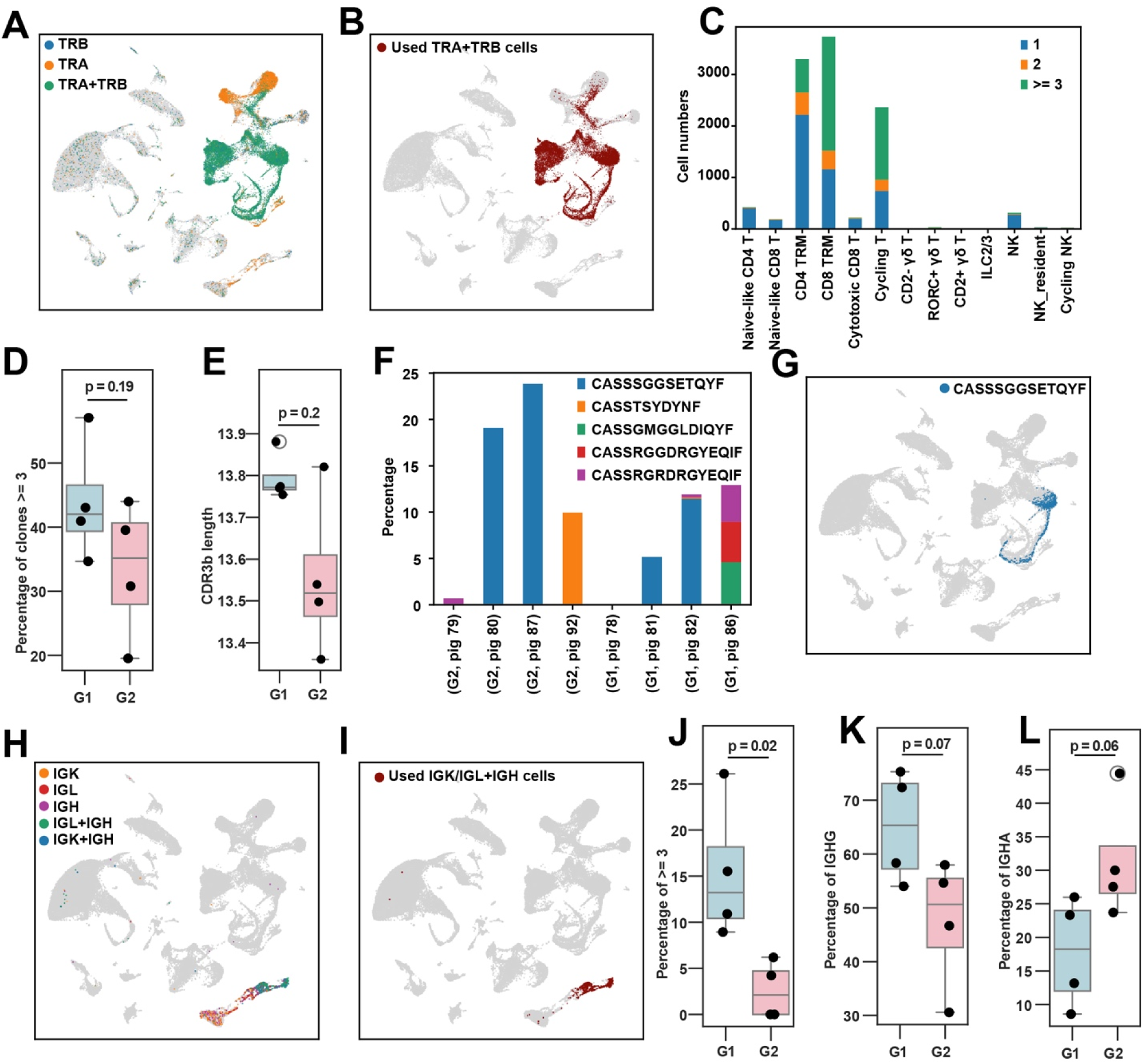
Immune receptor profiling of pulmonary αβ T and B cells. (A) UMAP plot showing the alignment of single TRA, single TRB, and paired TRA+TRB cells. (B) UMAP plot of T/NK/ILC cell clusters with paired TRA+TRB, which were used for downstream analysis. (C) Number of cells expressing 1, 2, or ≥3 clones identified by CDR3β sequence. (D) Box plot showing the proportion of cells expressing ≥3 clones across vaccinated groups (G1 and G2). Each dot represents an individual animal. Statistical significance was assessed by t-test. (E) Box plot showing the average length of CDR3β amino acid sequences by vaccinated groups (G1 and G2). Each dot represents the mean CDR3β length of a sample. Statistical significance was assessed by t-test. (F) Proportion of top 5 expanded CDR3β sequences in each animal. (G) Cells expressing the CDR3β sequence CASSSGGSETQYF. (H) UMAP plot showing the alignment of cells expressing single IGK, IGL, IGH, and paired IGL/IGK+IGH chains. (I) UMAP plot of B cells with paired IGL/IGK+IGH, which were used for downstream analysis. (J) Box plot of the proportion of cells expressing ≥3 clones (combined IGL/IGK and IGH CDR3) in vaccinated groups (G1 and G2). Each dot represents an individual animal. Significance was determined by t-test. (K– L) Box plots of the proportion of cells expressing *IGHG* (K) and *IGHA* (L) in vaccinated groups (G1 and G2). Each dot represents an individual animal. Significance was determined by t-test. G1: *CD1D−/−* vaccinated and challenged; G2: *CD1D−/+* vaccinated and challenged.

Next, we examined expanded clonotypes taking advantage of the fact that V(D)J recombination at the TCR and BCR loci can be used as endogenous barcodes to trace T and B cell clonotypes as they expand or transition through different states. The highest concentrations of clonally expanded T cells were among CD4^+^ TRMs, CD8^+^ TRMs, and cycling T cells (Figure 7C). *CD1D−/−* pigs had a higher frequency of expanded clones compared to *CD1D−/+* pigs (Figure 7D), whereas the *CD1D−/+* group had shorter average CDR3β lengths than the *CD1D−/−* group (Figure 7E), which has been associated with converging TCR motif signatures in a number of diseases (57, 58). Several of the most expanded clones were found in more than one pig, including pigs of both genotypes (Figure 7F). Indeed, the most expanded clonotype (CASSSGGSETQFY) was a vigorously proliferating CD8^+^ TRM, which was detected in four of the eight samples (Figure 7G). Interestingly, this sequence differed by a single amino acid from a curated human TCR CDR3β sequence (CASSSG<u>E</u>SETQYF) in the IEDB database (https://www.iedb.org/), that recognizes the IAV Matrix protein 1 epitope GILGFVFTL when presented by human HLA-A*0201 (59). It is notable that several expanded clones in this study were identified among expanded clones from a prior study which used lung samples collected from *CD1D−/−* and *CD1D−/+* pigs following infection with H1N1 pdmCA04 and subsequent re-challenge with a heterologous H1N1 virus (40) (Table S8), indicating their potential as IAV-reactive clonotypes.

A similar analysis of BCR chains was performed, with an average of 99% of paired IGK/IGL+IGH cells mapping to B cells (Figures 7H–I, Data file 2). We found that *CD1D−/−* pigs also had a higher frequency of expanded clones compared to *CD1D−/+* pigs (Figure 7J). Additionally, *CD1D−/−* B cells expressed a higher proportion of *IGHG* (Figure 7K) and a lower proportion of *IGHA* (Figure 7L) heavy chain transcripts compared to *CD1D−/+* B cells. Unlike in T cells, we did not identify expanded B cell clonotypes in more than one sample, due to the high diversity of heavy chain CDR3 sequences (Figure S3). Together, these findings indicate that NKT cell-derived stimuli influenced the immune receptor repertoire of the lung.

Finally, we profiled γ and δ TCR chain expression using newly developed primers that target the C regions of each chain (Table S1). Assembled V(D)J sequences were blasted against the international ImMunoGeneTics (IMGT) germline TRGV, TRGJ, and TRDJ databases (60). Because Vδ genes are not annotated in IMGT, Vδ sequences were assigned according to pig TRAV/TRDV sequences from our previous publication (61) (Data file 2). Over the eight samples, we obtained an average of 206,307,805 reads per TCR library with 70% of paired γ+δ cells mapping to γδ T cell clusters (clusters 11–13). An additional 21% mapped to the CD8^+^ TRM cluster (cluster 4), which we discovered contained a mixture of CD8[αβ and γδ T cells (Figure 8A). Only T cells with paired, productive γ and δ TCR chains were used for further analysis (Figure 8B). As expected, the γ and δ TCR repertoire was comprised of a limited set of VJ and C segments (Figure 8C). Using CDR3γ, we found that CD8^+^ TRM, CD2*^−^*, and *RORC*^+^ γδ T cell subsets had high proportions of expanded clonotypes, whereas CD2^+^ γδ T cells had a low proportion (Figure 8D). Notably, the five most common CDR3γ sequence, which were shared across samples, comprised a high fraction of each pig’s total sequences, ranging from between 20 and 42% (Figure 8E). In terms of γ chain V, J, and C segment usage, CD2^+^ γδ T cells were the most diverse, while CD2*^−^* γδ T cells were the least diverse (Figures S4A–C). In fact, CD2*^−^* and *RORC*^+^ γδ T cells were dominated by *TRGV3*, *TRGJ5*, and *TRGC5* segments (Figures 8F–H). We also found that CDR3γ lengths in the CD2*^−^* subset were greater and less variable than in the other γδ subsets (Figure S4D). As regards the δ chain, we found that the CD2^+^, CD2*^−^*, and *RORC*^+^ γδ subsets had greater V segment diversity than CD8^+^ TRM-resident γδ T cells (Figures S4E–H). We next compared *CD1D−/−* and *CD1D−/+* pigs to assess if NKT cell status affected the pulmonary γδ TCR repertoire. *CD1D−/+* pigs tended to have a higher frequency of expanded clones and lower repertoire diversity compared to *CD1D−/−* pigs (Figures 8I and S4I). Other differences included that the CAGWNYSSRWIKIF clonotype was much more prevalent in *CD1D−/+* pigs and that *CD1D−/−* pigs exhibited greater overlap in CDR3γ sequences and more convergent V and J segment usage (Figures 8E and S4J–K), indicating a more homogeneous γδ TCR repertoire in *CD1D−/−* pigs.

**Figure 8.**
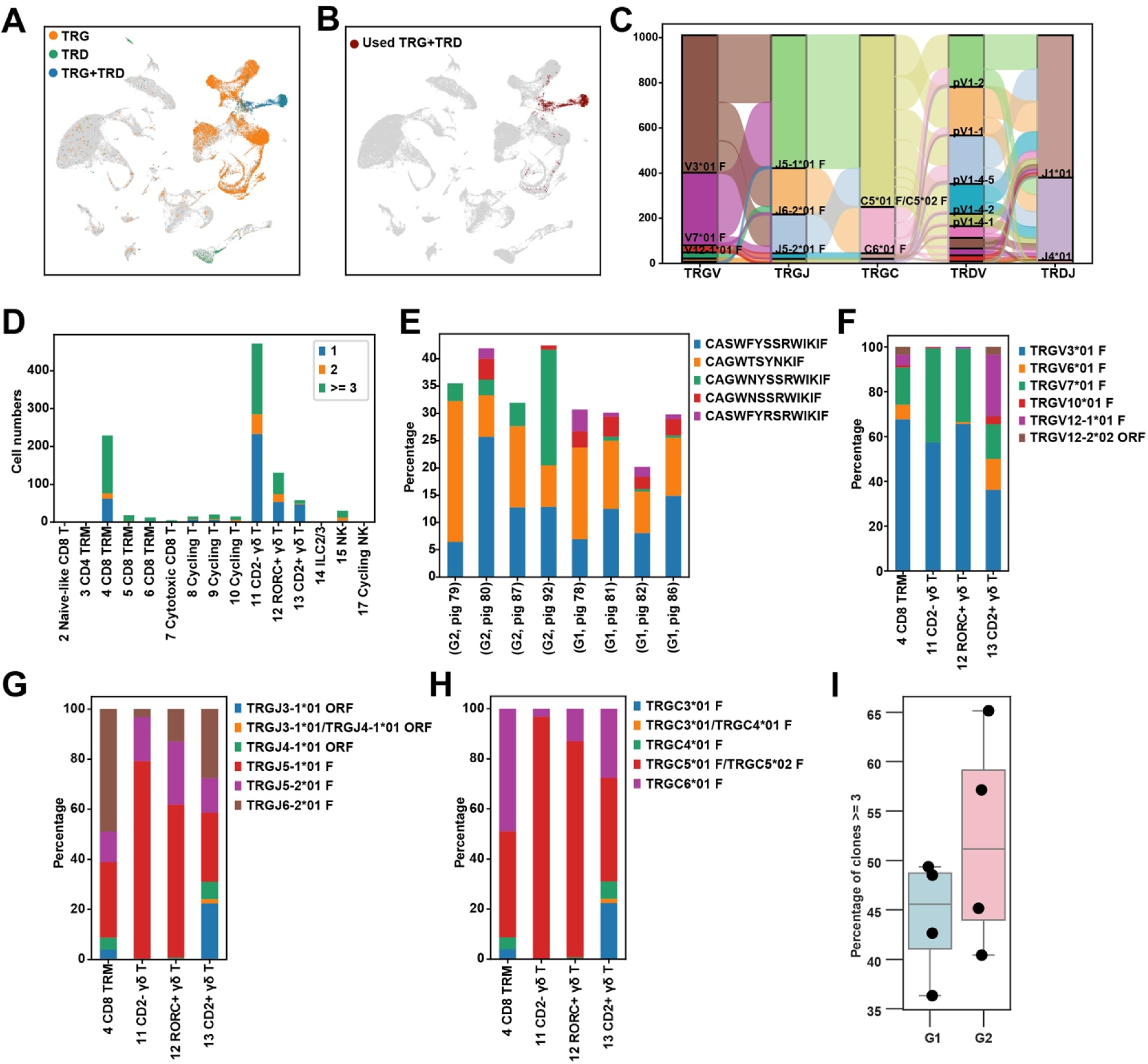
Immune receptor profiling of pulmonary γδ T cells. (A) UMAP plot showing the alignment of single TRG, single TRD, and paired TRG+TRD cells. (B) UMAP plot of cells with paired TRG+TRD, which were used for downstream analysis. (C) Sankey-plot showing the number of cells expressing TRG and TRD VJC gene segments, as well as their recombination patterns. (D) Number of cells expressing 1, 2, or ≥3 clones identified by CDR3γ sequence. (E) Proportion of top 5 expanded CDR3γ sequences in each animal. (F–H) Proportion of cells expressing *TRGV* (F), *TRGJ* (G), *TRGC* (H) segments by cell type. (I) Box plot of the proportion of cells expressing ≥3 clones in G1 and G2 groups. Each dot represents an individual animal. G1: *CD1D−/−* vaccinated and challenged; G2: *CD1D−/+* vaccinated and challenged.

### NKT cells modulate virus-specific T cell kinetics

Finally, interferon-γ (IFN-γ) ELISpot assays were performed to quantify virus-reactive cells in PBMCs and lung cell suspensions. Unvaccinated pigs did not develop measurable TX98- or CO99-specific IFN-γ producing PBMCs until 5 DPC, when they were detected at low levels. On the other hand, both vaccinated groups developed TX98- and CO99-reactive cells by 14 DPV, and their concentration increased substantially after challenge (Figures 9A–B). Interestingly, T cell kinetics differed by genotype; at 14 DPV vaccinated *CD1D−/+* pigs had more TX98- and CO99-reactive PBMCs than *CD1D−/−* pigs; at 20 DPV the groups were comparable; but at 5 DPC, *CD1D−/−* pigs had significantly more virus-reactive cells than *CD1D−/+* pigs. The same pattern was observed in 5 DPC lung cells (Figures 9C–D). We also analyzed PBMCs using an IL-2 ELISpot assay, which presented similar results to the IFN-γ ELISpot assay, except that there was no significant difference between genotypes at 14 DPV (Figures 9E– F). Collectively, these results indicate that NKT cells alter the kinetics of the cellular response, with virus-reactive cells in *CD1D−/−* pigs taking longer to develop but eventually reaching higher concentrations than in *CD1D−/+* pigs. This agrees with our single-cell data showing that *CD1D−/−* lung samples had higher frequencies of expanded αβ T cell clones than *CD1D−/+* lungs (Figure 7D).

**Figure 9.**
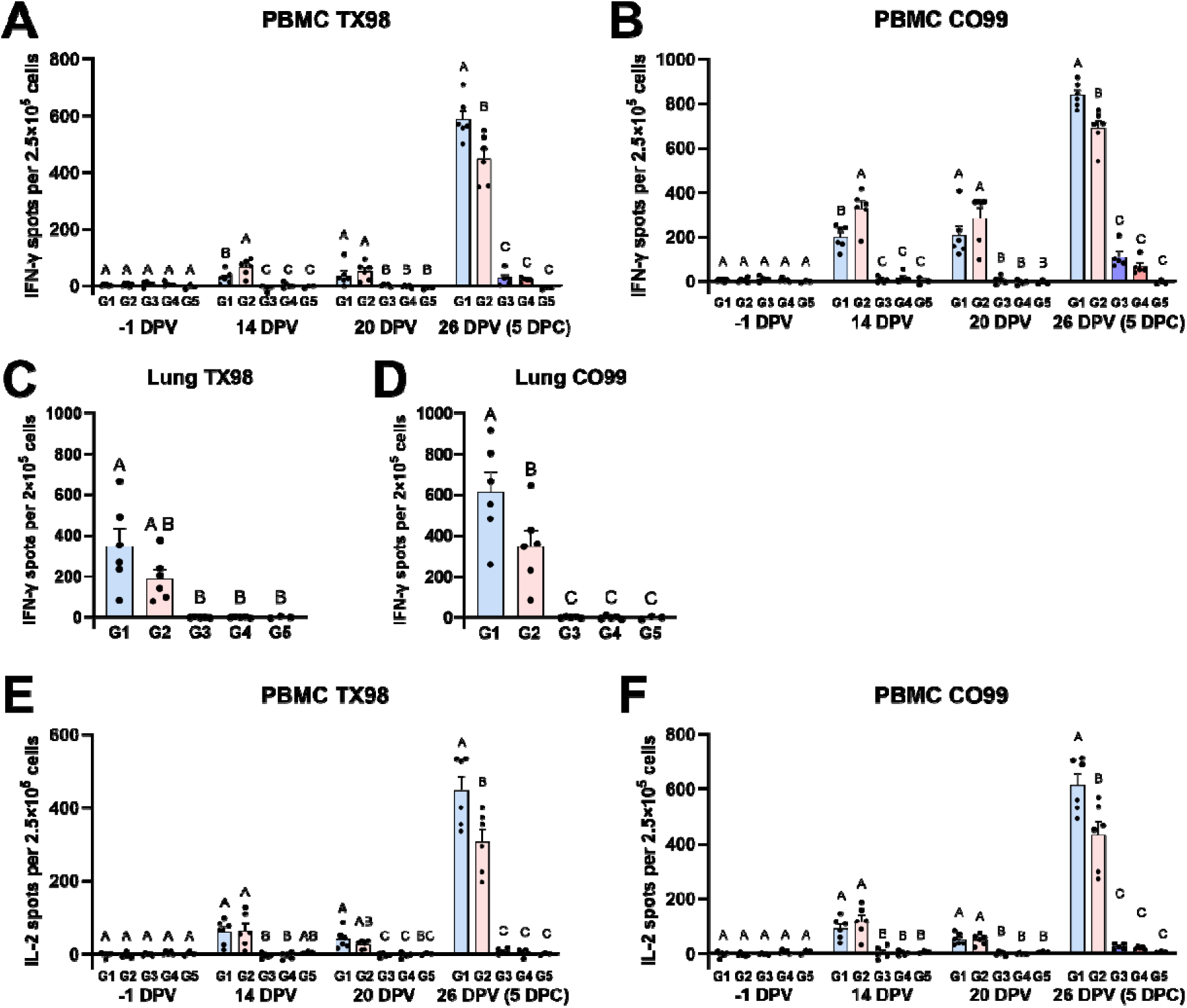
Cellular responses to TX98 and CO99 measured by ELISpot assays. (A, B) Interferon-γ (IFN-γ production by PBMCs collected at -1, 14, 20, and 26 DPV (5 DPC) after incubation with live TX98 (A) and CO99 (B) virus. (C, D) IFN-γ production by lung cells collected at 5 DPC after incubation with live TX98 (E) and CO99 (F) virus. (E, F) Interleukin-2 (IL-2) production by the same PBMCs after incubation with live TX98 (C) and CO99 (D) virus. A statistically significant difference (*p* < 0.05) between two groups is indicated by different letters. Data are represented as mean ± SEM. Symbols represent individual pigs. G1: *CD1D−/−* vaccinated and challenged; G2: *CD1D−/+* vaccinated and challenged; G3: *CD1D−/−* not vaccinated and challenged; G4: *CD1D−/+* not vaccinated and challenged; G5: *CD1D−/+* not vaccinated, not challenged.

## Discussion

The current work assessed the contribution of pig NKT cells to immunity induced by a MLV vaccine with a truncated NS1 protein, which in swine generally affords robust protection against heterologous IAV infection. Our premise was based on previous reports that NKT cells function as a type of universal T helper cell enhancing both humoral and cellular immune responses, especially in mucosal tissues like the lower respiratory tract (lungs) where they are enriched (4). This is supported by the fact that compared to standard mice, NKT cell-deficient mice develop weaker cellular and humoral responses after IAV immunization (10, 11) While swine possess the NKT cell-CD1d system, the concentration of NKT cells in most pigs is considerably lower than in most inbred mouse strains (37, 62). Moreover, the functional diversity of pig NKT cells differs considerably from the NKT0/1/2/17 subset differentiation paradigm established in mice (63). Nevertheless, porcine NKT cells react strongly to α-GalCer *in vitro* and *in vivo* (37, 45, 62), and pigs co-administered α-GalCer in combination with inactivated IAV vaccines develop greater humoral and cellular immune responses than pigs administered vaccine alone (3). In fact, α- GalCer is capable of triggering vaccine associated enhanced respiratory disease in pigs, an immunopathological condition where vaccination causes enhanced lung disease rather than protection after a subsequent infection (64).

The current work found no evidence that vaccination was less effective at decreasing viral clearance or reducing pulmonary pathology in NKT cell-deficient pigs compared to NKT cell-intact pigs. This aligns with a previous mouse study which found no difference in virus titers, pulmonary pathology, or mortality between IAV-infected *CD1d*-deficient and standard mice that had been previously immunized with a sublethal dose of live virus (13).

In terms of immunological parameters, *CD1D−/−* and *CD1D−/+* pigs developed similar antigen-specific humoral responses, except that *CD1D−/+* pigs skewed towards a higher HA-specific IgG1/IgG2 ratio.

This resembles prior mouse studies showing that NKT cell activation skews the humoral response towards a Th2-driven IgG1 response (65). While the mouse IgG subclass-cytokine paradigm may not directly apply to pigs, our findings suggest that NKT cells did influence the underlying T helper response.

The impact on cellular responses was more pronounced, with virus-specific cells in *CD1D−/−* pigs taking longer to develop but eventually surpassing *CD1D−/+* pigs, in both blood and lung tissue. This observation may be related to the fact that, because NKT cells are capable of sequentially secreting pro- and ant-inflammatory cytokines, they often play Janus-like opposing roles in different phases of the immune response (66). In this regard, our observations support that NKT cells played a stimulatory role during the initial induction of virus-reactive T cells following vaccination but then switched to a suppressive role once pigs became infected with live virus, perhaps as a mechanism to prevent exacerbated inflammation. Our observation agrees with a prior study which showed that virus-specific CD8^+^ T cell responses were significantly greater in *CD1D−/−* compared to standard mice after vaccination with an inactivated IAV vaccine (12). The authors attributed this suppression to NKT cell-dependent IDO expression. We found it interesting that NKT cell status seemed to affect the distribution of T cells in infected lungs, in that without them, CD3^+^ T cells did not accumulate around airways as they did in NKT cell-sufficient pigs. This observation is reminiscent of a previous study in which we found that pigs vaccinated with α-GalCer and an inactivated IAV vaccine had a higher density of intra-epithelial CD3^+^ T cells associated with the bronchiolar epithelium than did vaccinated pigs that did not receive α-GalCer (64). Together, these results indicate that NKT cells play an important role in coordinating cellular responses following IAV encounter, including diminishing the amount of virus-specific T cells that accumulate around smaller-caliber airways in the infected lung.

Our single-cell transcriptomics analysis revealed that ablating NKT cells changed the cellular composition of the lung at 5 DPC, including that *CD1D−/−* pigs were enriched for TRMs, consistent with the ELISpot and flow cytometry results. It also revealed that NKT cell deletion downregulated a substantial number of genes and pathways in TRMs and other lymphocyte populations, several of which are integral to lymphocyte trafficking, development, proliferation, differentiation, and effector functions. While this seems contradictory given that *CD1D−/−* pigs had higher TRM levels than *CD1D−/+* pigs, lymphocyte homeostasis is governed by a balance of activation and inhibitory signals. Thus, our results may indicate negative feedback mechanisms involving TCR/BCR signaling, inhibitory receptors, and cytokines following activation in *CD1D−/−* lymphocytes. Also notable is that *CD1D−/−* lung macrophages and mono-mac cells upregulated pathways associated with anti-inflammatory responses and tissue repair. This suggests that NKT cells activated in response to influenza vaccination/infection play a role in controlling the functional program of macrophages. This is relevant as lung macrophages are essential players in shaping the outcome of immune responses and disease outcomes in viral respiratory infections (67, 68).

Our single-cell TCR and BCR sequencing analysis found that *CD1D−/−* pigs had higher numbers of expanded αβ T cell clones in their lungs, which agrees with our ELISpot assays showing more virus-specific T cells in *CD1D−/−* than *CD1D−/+* pigs after infection. Most of the expanded clones were within CD4^+^ TRM and CD8^+^ TRM clusters, consistent with the notion that these cells contain antigen experienced T cells capable of rapidly responding to IAV infection. Interestingly, several of our most expanded sequences aligned exactly to CDR3β sequences from a prior study where we performed single cell TCR sequencing on lung T cells from *CD1D−/−* than *CD1D−/+* pigs sequentially infected with two heterologous H1N1 IAVs (40). Thus, we may have uncovered clonotypes capable of identifying the same IAV antigens. Among the most expanded clonotypes, we identified a clone with a CDR3β sequence almost identical to a human motif that recognizes an immunodominant epitope from Matrix protein 1 (69, 70). This may arise because peptide binding motifs of some common swine leukocyte antigen (SLA) molecules partly overlap with the binding motifs of human HLA molecules (71).

Our scBCRseq analysis found that *CD1D−/−* pigs also had higher numbers of expanded lung B cell clones compared to *CD1D−/+* pigs, indicating that NKT cells may also play a role in modulating B cell expansion. Additionally, there was also a shift in heavy chain usage, with *CD1D−/−* B cells favoring IgG and *CD1D−/+* B cells favoring IgA transcripts. This could be related to the cognate or non-cognate help that NKT cells are known to provide to B cells, which can affect immunoglobulin heavy chain usage (6–9).

Our novel γδ TCR assay produced several interesting findings. As in humans, there was greater clonal diversity in the pool or rearranged *TRD* genes than *TRG* genes, a small number of highly expanded clones accounted for a high proportion of all *TRG* sequences, and the TRG repertoire was comprised of a high proportion of public sequences whereas the *TRD* repertoire is mostly private (72). We also found that the Th17-associated CD2^−^ and *RORC*^+^ subsets had lower diversity in γ chain V-J and C segment usage and much higher proportions of expanded clonotypes compared to the CD2^+^ subset. Moreover, both CD2^−^ and *RORC*^+^ γδ T cells were dominated by *TRGV3*, *TRGJ5*, and *TRGC5* segments, indicating that they share a common ancestor. The CDR3γ sequences of CD2^−^ γδ T cells were longer but more uniform than the other subsets. This agrees with their high level of clonal expansion as T cell proliferation often causes the resulting TCR repertoire to become more focused (73). We also identified a population of γδ T cells within the CD8^+^ TRM cluster that had a TRG VJC segment usage pattern similar to the CD2^+^ γδ T cells. Like the CD8^+^ αβ T cells within this cluster, these γδ T cells were enriched in memory- and cytotoxicity-associated genes. Regarding genotype differences, our observation that *CD1D−/+* pigs tended to have a higher frequency of expanded clones and a lower TCR repertoire diversity compared to *CD1D−/−* pigs suggests that NKT cells influence pulmonary γδ T cell differentiation. However, additional studies are required to corroborate this.

Also of interest was the response of unvaccinated naïve *CD1D−/−* and *CD1D−/+* pigs to IAV infection since several studies have shown that in mice NKT cells contribute to early-innate IAV responses that inhibit virus replication and protect against disease (28–31). Although not significant, we found that virus titers in *CD1D−/+* pigs tended to be higher than *CD1D−/−* pigs, indicating that in pigs, NKT cell responses do not inhibit, and may even support virus replication. Indeed, we have observed, in several prior IAV challenge studies using pandemic H1N1 (CA04), that virus shedding is somewhat delayed in naïve *CD1D−/−* compared to *CD1D−/+* pigs (Figure S5). This discrepancy from mice raises a cautionary note about interpreting the results of NKT cell studies conducted in animal models since we are not yet certain which model best represents human NKT cell biology.

In conclusion, we found that genetically ablating NKT cells did not substantially alter virus load or pulmonary pathology following IAV infection in pigs, regardless of their vaccination status. Nonetheless, removing NKT cells significantly altered the nature of the developing immune response, particularly as regards the dynamics of virus-specific T cell accumulation and the localization of T cells in the lung after infection. In this regard, our data supports that NKT cells help to constrain the accumulation of virus-specific T cells in the lung, perhaps as a mechanism to prevent excessive pulmonary inflammation. It stands to reason that, although these changes did not affect disease in the current investigation, they are likely to be of significance in other settings, such as TRM persistence in the respiratory tract and immunity against heterosubtypic viruses. The same may also be true in humans and therefore deserves further study.

## Acknowledgments

Funding for this study was provided through grants from the National Institutes of Health grant AI158477 and the U.S. Department of Agriculture grant 2021-67015. Additional funding was provided by the National Bio and Agro-Defense Facility (NBAF) Transition Fund from the State of Kansas, the AMP and MCB Core of the Center on Emerging and Zoonotic Infectious Diseases (CEZID) of the National Institutes of General Medical Sciences under award number P20GM130448, and the NIAID supported Center of Excellence for Influenza Research and Response (CEIRR) under contract number 75N93021C00016. The funding for the National Swine Resource and Research Center is from the National Institute of Allergy and Infectious Disease, the National Institute of Heart, Lung and Blood, and the Office of the Director (U42OD011140). LSU acknowledges support from an Institutional Development Award (IDeA) from the National Institute of General Medical Sciences of the National Institutes of Health under grant number P20GM130555-5011 (to MC), U.S. Department of Agriculture’s (USDA) National Institute of Food and Agriculture (NIFA) Agriculture and Food Research Initiative and American Rescue Plan Act through USDA Animal and Plant Health Inspection Service (APHIS) competitive grant number 2023-70432-39465 (to MC) and the School of Veterinary Medicine, Louisiana State University (PG009641) (to MC). We gratefully thank Shristi Ghimire, Shanmugasundaram Elango, and Sujan Kafle at Kansas State University for technical assistance and Melissa Samuel, Kristin Whitworth, and Anna Spate at University of Missouri for husbandry, maintenance, and genotyping of the *CD1D* modified pigs. We also thank members of the Louisiana Animal Disease Diagnostic Laboratory for their assistance in histological processing of tissues, slide preparation and staining.

