## Supplemental Figures for "Pigs lacking Natural Killer T cells have altered cellular responses to influenza"

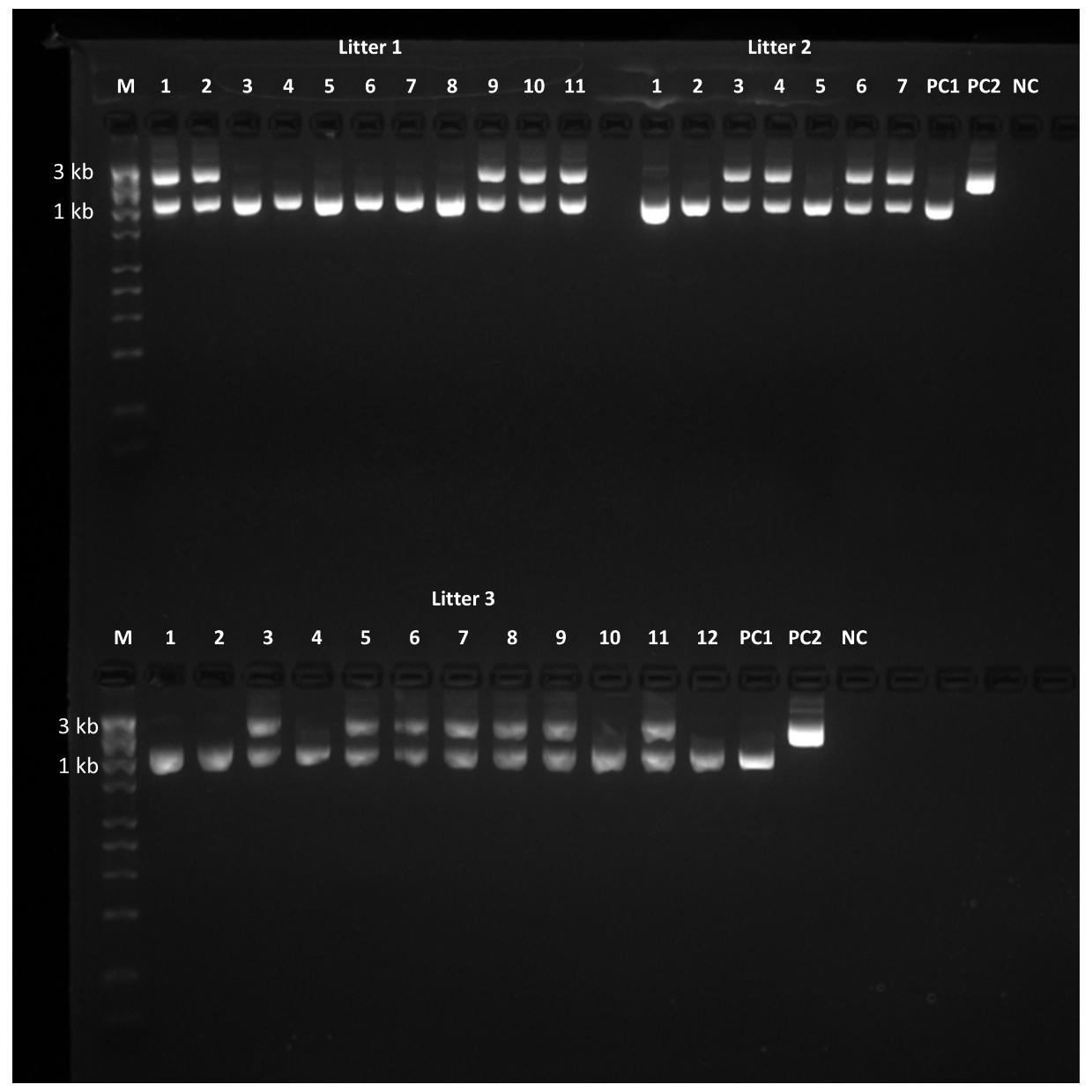

**Figure S1.** *CD1D* genotyping results. *CD1D* genotype of pigs from three litters was confirmed by PCR targeting a 2,788 bp product of the endogenous porcine *CD1D* gene. Pigs that possessed an edited *CD1D* allele produced a deletion of 1,598 bp resulting in a modified product of 1,189 bp. A single 1,189 bp band was detected in homozygous (*CD1D−/−*) pigs, whereas two PCR products at 1,189 and 2,787 bp were detected in heterozygous (*CD1D−/+)* pigs. M: DNA molecular marker; PC1: a positive control for the modified allele; PC2: a positive control for an unmodified, “wildtype” sample; NC: negative control.

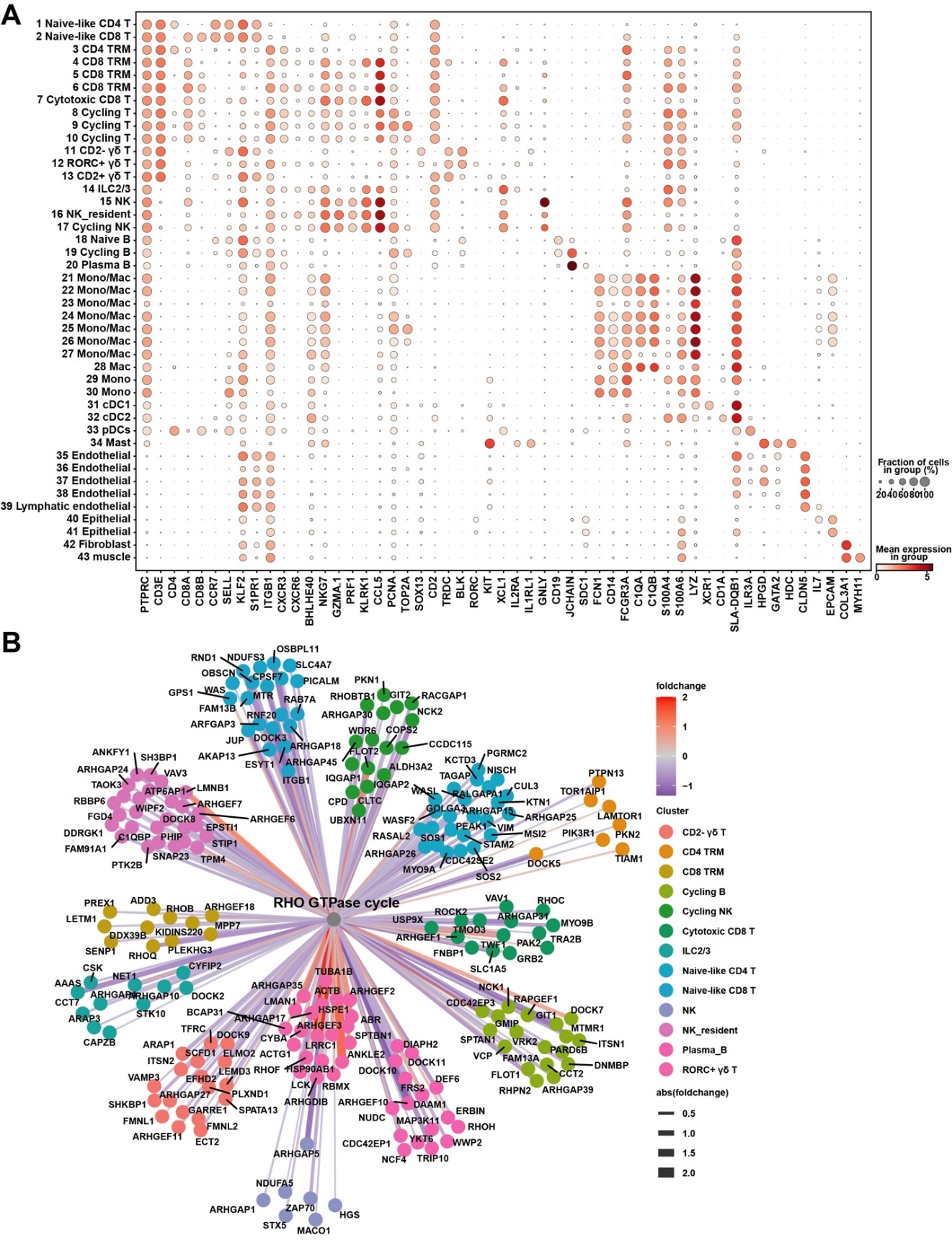

**Figure S2.** (A) Dot plot showing the mean expression of selected marker genes in each cluster from Figure 5A. (B) Network graph showing cluster-specific DEGs involved in the RHO GTPase cycle pathway. Edge color indicates positive or negative fold change in G1 compared to G2, and edge thickness represents the absolute fold change. G1: *CD1D−/−* vaccinated and challenged; G2: *CD1D−/+* vaccinated and challenged.

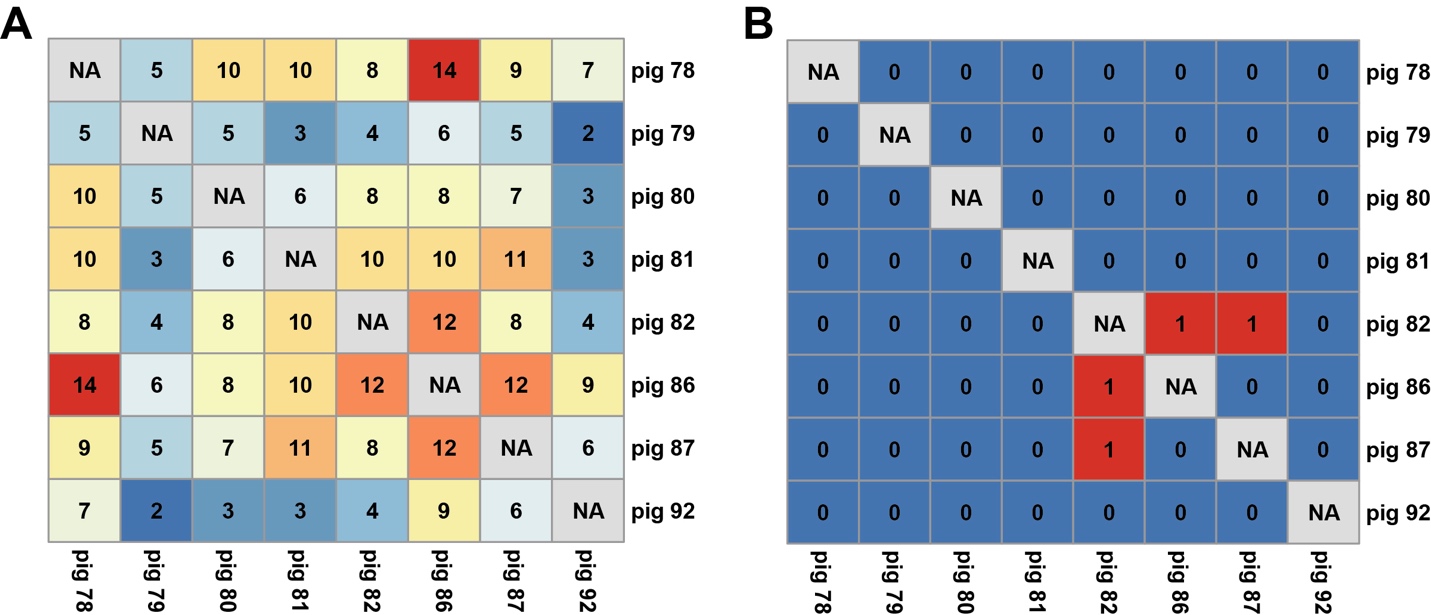

**Figure S3.** Heatmaps showing the overlap in cell numbers across samples for BCR light chain CDR3s (A) and heavy chain CDR3s (B).

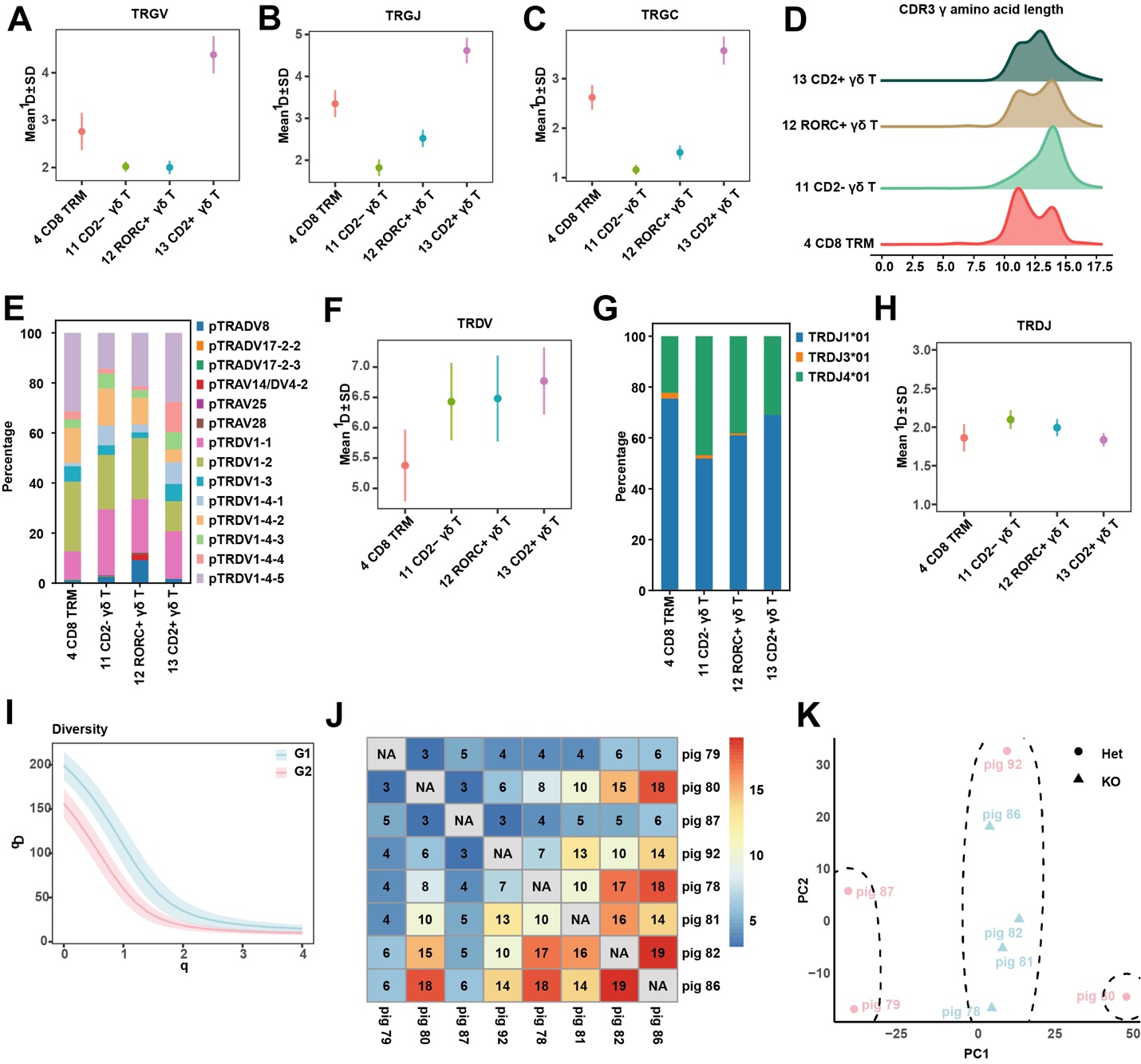

**Figure S4**. (A–C, F, H) VJC segment diversity measured using the Shannon–Wiener index, corresponding to the Hill diversity index at order q = 1 for each cell type. (D) CDR3γ amino acid length distribution across cell types. (E, G) Proportion of cells expressing TRDV (E) and TRDJ (G) segments by cell type. (I) CDR3γ diversity across varying Hill diversity orders. (J) Heatmaps showing the number of overlapping CDR3γ across samples. (K) Principal component analysis (PCA) of TRG and TRD VJC segment usages by sample.

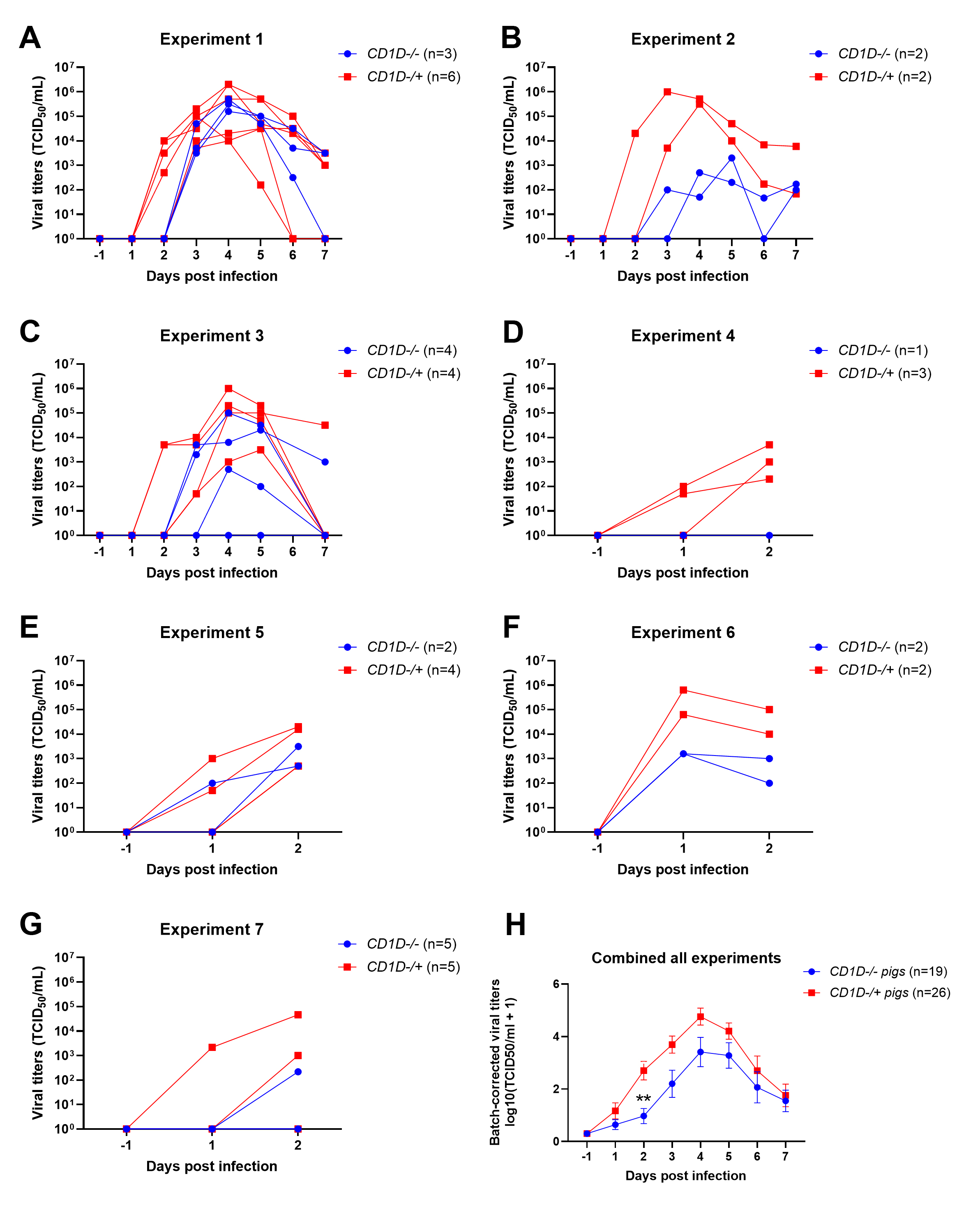

**Figure S5**. (A–G) Viral titers in nasal swabs from seven prior experiments where *CD1D−/−* and *CD1D−/+* pigs between 4 and 6 weeks of age were intratracheally infected with pandemic H1N1 A/California/04/2009 influenza A virus. Each line represents an individual pig. (H) Mean viral titers across all experiments are presented as mean ± SEM. To minimize batch effects, titers were log₁₀-transformed and normalized by centering each batch to the overall mean (adjusted value = raw − batch mean + grand mean). The adjusted values were used for statistical analyses. Treatment and time effects were assessed using a mixed-effects model (REML), followed by Sidak’s multiple comparisons test for pairwise comparisons.
