## Supplemental Table for "Pigs lacking Natural Killer T cells have altered cellular responses to influenza"

| Primer name | Target | 5'-3' Sequence | Accession | Ref. type |
| --- | --- | --- | --- | --- |
| TCRa outer | TRAC | ATCGGTGCTTTTGCTCCAAG | MN086839.1 | mRNA |
| TCRa inner | TRAC | GTGGGCTCCGAGTCTTTTGT | MN086839.1 | mRNA |
| TCRb outer | TRBC | TCAGACAGTAGCTGGAGTCATTGAG | AB079894.1 | DNA |
| TCRb inner | TRBC | TCCGATGGTTCAAACACGGC | AB079894.1 | DNA |
| TCRg inner | TRGC3/4/6 | TCCAGAAGACAAAGGTATGTTCCA | AB185445.1 | mRNA |
| TCRg inner | TRGC5 | TCAAGAAGACAAAGATGTGTCCCA | BK074883 | DNA |
| TCRd outer | TRDC | CTCCATACTGACCAAGCTTGACGG | AB182371.1 | DNA |
| TCRd inner | TRDC | GACCACGATAGCAGGGTCATAT | AB182371.1 | DNA |
| IgA outer | IGHAC | TGCACTTGGCACTTCAGGAT | AB699688.1 | DNA |
| IgA inner | IGHAC | CAATAACGCCCTCGCGACTA | AB699688.1 | DNA |
| IgG outer | IGHGC | CTGAGGGAGTAGAGCCCTGA | AB699686.1 | DNA |
| IgG inner | IGHGC | GCTCGGGGAAGTAGCTTGAG | AB699686.1 | DNA |
| IgM/D outer | IGH(M/D)C | AAGTACTTGCCGCCTCTCAG | AB699686.1 | DNA |
| IgM/D inner | IGH(M/D)C | GATGTTCTGGCTGCTGACCT | AB699686.1 | DNA |
| Kappa outer | IGKC | GAAGCTTTTGACCAGAGGGGA | KF561240.1 | mRNA |
| Kappa inner | IGKC | TCCAGGATGCCACTGCTTTG | KF561240.1 | mRNA |
| Lambda outer | IGLC | CGTCACTGTCTTCTCCACAATG | M59322.1 | mRNA |
| Lambda inner | IGLC | TCTGTTTCGAGGGCTTGGTG | M59322.1 | mRNA |

Table S1. Porcine custom primer sets for scTCR/BCRseq

Table S2. Frequency (mean ± SEM) of leukocyte populations in lungs at 5 days post challenge

| Immune cell population | Group 1: Vaccinated *CD1D−/−* | Group 2: Vaccinated *CD1D−/+* | Group 3: Unvaccinated *CD1D−/−* | Group 4: Unvaccinated *CD1D−/+* | Group 5: Negative *CD1D−/+* ^a^ |
| --- | --- | --- | --- | --- | --- |
| CD3^+^ (of lymphocytes) | 47.3 ± 3.9 | 35.3 ± 4.1 | 29.1 ± 2 | 26.8 ± 1.2 | 21.5 ± 1.7 |
| αβ cells (CD3^+^TCRδ^-^ of lymphocytes) | 35.1 ± 3.3 | 27 ± 2.7 | 16.7 ± 1.3 | 15.7 ± 1.5 | 14.5 ± 1.9 |
| γδ cells (CD3^+^TCRδ^+^ of lymphocytes) | 7.6 ± 1 | 4 ± 0.6 | 8.9 ± 1 | 7.8 ± 0.7 | 5.4 ± 2.3 |
| CD4^-^CD8α^+^ (of CD3^+^) | 48.5 ± 2.9 | 48.6 ± 2.1 | 42.5 ± 4 | 41.2 ± 1.9 | 37 ± 5.6 |
| CD4^+^CD8α^+^ (of CD3^+^) | 24.2 ± 2.1 | 34.1 ± 3.4 | 23.4 ± 2.3 | 23.3 ± 1.4 | 28.5 ± 3.6 |
| CD4^+^CD8α^-^ (of CD3^+^) | 16.3 ± 0.9 | 12.6 ± 1.2 | 13.1 ± 0.7 | 13.9 ± 2.2 | 16.9 ± 2.1 |
| CD8α^+^ CD8β^+^ (of CD3^+^) | 40.8 ± 4.1 | 39.9 ± 1.9 | 23.3 ± 2.9 | 24.6 ± 2 | 20.2 ± 4.3 |
| NK cells (CD8α^+^CD3^-^ of lymphocytes) | 11.1 ± 1.4 | 12.9 ± 1.2 | 15.4 ± 0.8 | 18.1 ± 1.9 | 13 ± 0.9 |
| Macrophages (CD14^+^CD11b^-^CD163^+^ of leukocytes) | 15.8 ± 0.8 | 19.3 ± 1.7 | 20.1 ± 2.9 | 17.1 ± 1.3 | 19.3 ± 1.8 |
| Monocytes (CD14^+^CD11b^-^CD163^-^ of leukocytes) | 13.9 ± 1.5 | 20.7 ± 3.4 | 15.2 ± 1.3 | 19.8 ± 1.3 | 23.1 ± 0.7 |
| Neutrophils (CD14^+^CD16^+^CD163^-^ of leukocytes) | 2.4 ± 0.3 | 3.8 ± 0.6 | 2.1 ± 0.2 | 2.3 ± 0.3 | 3.5 ± 0.2 |

^a^ The lung of negative *CD1D*-/+ pigs were collected at 17 days post vaccination.

Table S3. Frequency (mean ± SEM) of leukocyte populations in tracheobronchial lymph nodes at 5 days post challenge

| Immune cell population | Group 1: Vaccinated *CD1D−/−* | Group 2: Vaccinated *CD1D−/+* | Group 3: Unvaccinated *CD1D−/−* | Group 4: Unvaccinated *CD1D−/+* | Group 5: Negative *CD1D−/+* ^a^ |
| --- | --- | --- | --- | --- | --- |
| CD3^+^ (of lymphocytes) | 56.6 ± 5.1 | 62.9 ± 6.5 | 20.7 ± 0.8 | 30.1 ± 5.7 | 68 ± 6.7 |
| αβ cells (CD3^+^TCRδ^-^ of lymphocytes) | 50.6 ± 4.8 | 60.2 ± 6.7 | 20.4 ± 1 | 28.2 ± 5.1 | 62.2 ± 6.4 |
| γδ cells (CD3^+^TCRδ^+^ of lymphocytes) | 2.5 ± 0.1 | 1.8 ± 0.1 | 1.5 ± 0.2 | 2.1 ± 0.2 | 3.5 ± 0.7 |
| CD4^-^CD8α^+^ (of CD3^+^) | 21.9 ± 1 | 25.5 ± 1.6 | 23.2 ± 1.3 | 26.8 ± 2.5 | 25.3 ± 1.5 |
| CD4^+^CD8α^+^ (of CD3^+^) | 37.8 ± 6.2 | 46.1 ± 4.4 | 28.4 ± 5.5 | 23 ± 2.8 | 28.1 ± 7.2 |
| CD4^+^CD8α^-^ (of CD3^+^) | 36.1 ± 5.9 | 25.7 ± 3.2 | 37.4 ± 5.3 | 38.6 ± 4.9 | 42.1 ± 6.8 |
| CD8α^+^ CD8β^+^ (of CD3^+^) | 20.6 ± 1.1 | 21.9 ± 1.3 | 22.3 ± 2.2 | 23.8 ± 1.7 | 20.1 ± 0.5 |
| NK cells (CD8α^+^CD3^-^ of lymphocytes) | 10.3 ± 1.8 | 16.2 ± 4.2 | 9.5 ± 3.1 | 4.7 ± 1.6 | 9.1 ± 2.6 |
| Macrophages (CD14^+^CD11b^-^CD163^+^ of leukocytes) | 0.2 ± 0 | 0.6 ± 0.1 | 0.4 ± 0.1 | 0.4 ± 0.1 | 0.9 ± 0.2 |
| Monocytes (CD14^+^CD11b^-^CD163^-^ of leukocytes) | 0.6 ± 0.1 | 0.9 ± 0.2 | 1.1 ± 0.2 | 1 ± 0.1 | 1.7 ± 0.4 |
| Neutrophils (CD14^+^CD16^+^CD163^-^ of leukocytes) | 0.2 ± 0 | 0.4 ± 0.1 | 0.2 ± 0 | 0.3 ± 0.1 | 0.9 ± 0.6 |

^a^ The tracheobronchial lymph node of negative *CD1D*-/+ pigs were collected at 17 days post vaccination.

Table S4. Frequency (mean ± SEM) of leukocyte populations in blood at -1 days post vaccination

| Immune cell population | Group 1: Vaccinated *CD1D−/−* | Group 2: Vaccinated *CD1D−/+* | Group 3: Unvaccinated *CD1D−/−* | Group 4: Unvaccinated *CD1D−/+* | Group 5: Negative *CD1D−/+* |
| --- | --- | --- | --- | --- | --- |
| CD3^+^ (of lymphocytes) | 68.6 ± 3.4 | 68.8 ± 3.1 | 71.6 ± 3.1 | 68.1 ± 2.8 | 63.4 ± 2.9 |
| αβ cells (CD3^+^TCRδ^-^ of lymphocytes) | 30.9 ± 3.6 | 30.5 ± 2.4 | 35.4 ± 0.9 | 28.5 ± 1.9 | 28.9 ± 1 |
| γδ cells (CD3^+^TCRδ^+^ of lymphocytes) | 32.8 ± 4.7 | 30.4 ± 1.9 | 26.7 ± 3.4 | 29.2 ± 3.1 | 25.9 ± 5.7 |
| CD4^-^CD8α^+^ (of CD3^+^) | 17.3 ± 2.7 | 17.5 ± 1.1 | 18 ± 0.9 | 17.7 ± 0.9 | 22.1 ± 1.8 |
| CD4^+^CD8α^+^ (of CD3^+^) | 20.6 ± 8.1 | 13.5 ± 1.6 | 13.2 ± 2 | 11.6 ± 1.3 | 12.6 ± 3.4 |
| CD4^+^CD8α^-^ (of CD3^+^) | 32.2 ± 3.9 | 31 ± 2.1 | 39 ± 1.9 | 32.5 ± 2.9 | 33.4 ± 0.7 |
| CD8α^+^ CD8β^+^ (of CD3^+^) | 12.8 ± 2.1 | 11.7 ± 1.4 | 13.6 ± 0.9 | 12.6 ± 0.7 | 15.2 ± 2.1 |
| NK cells (CD8α^+^CD3^-^ of lymphocytes) | 6 ± 1.5 | 5.3 ± 1.8 | 3.3 ± 0.7 | 4.5 ± 0.8 | 5.3 ± 0.9 |
| Macrophages (CD14^+^CD11b^-^CD163^+^ of leukocytes) | 3.1 ± 0.4 | 3.4 ± 0.4 | 3.1 ± 0.4 | 3.1 ± 0.3 | 2 ± 0.3 |
| Monocytes (CD14^+^CD11b^-^CD163^-^ of leukocytes) | 36.5 ± 8.2 | 31.9 ± 5 | 31.4 ± 2.9 | 34.3 ± 2.8 | 47.8 ± 2.8 |
| Neutrophils (CD14^+^CD16^+^CD163^-^ of leukocytes) | 17.8 ± 1.5 | 12.7 ± 1.4 | 14.6 ± 2.9 | 11 ± 1.9 | 14.1 ± 2.3 |

Table S5. Frequency (mean ± SEM) of leukocyte populations in blood at 14 days post vaccination

| Immune cell population | Group 1: Vaccinated *CD1D−/−* | Group 2: Vaccinated *CD1D−/+* | Group 3: Unvaccinated *CD1D−/−* | Group 4: Unvaccinated *CD1D−/+* | Group 5: Negative *CD1D−/+* |
| --- | --- | --- | --- | --- | --- |
| CD3^+^ (of lymphocytes) | 71.9 ± 1.5 | 66.9 ± 4 | 61.9 ± 4.3 | 56.6 ± 3.3 | 51.5 ± 5.4 |
| αβ cells (CD3^+^TCRδ^-^ of lymphocytes) | 24.9 ± 2.4 | 27.3 ± 2.9 | 28 ± 1.5 | 25.9 ± 1.7 | 26.4 ± 7.4 |
| γδ cells (CD3^+^TCRδ^+^ of lymphocytes) | 33.9 ± 3.8 | 31.1 ± 2.1 | 26 ± 3.4 | 22.3 ± 3.4 | 17.4 ± 8.2 |
| CD4^-^CD8α^+^ (of CD3^+^) | 16.4 ± 1.3 | 17.5 ± 1 | 18.7 ± 2.2 | 18.7 ± 1.4 | 19.8 ± 3 |
| CD4^+^CD8α^+^ (of CD3^+^) | 14 ± 0.9 | 15.8 ± 1.3 | 15.5 ± 2.5 | 14.7 ± 2.3 | 23.9 ± 9 |
| CD4^+^CD8α^-^ (of CD3^+^) | 23.5 ± 2.6 | 23.3 ± 2.3 | 29.2 ± 1.5 | 29.9 ± 2.9 | 30.2 ± 8.9 |
| CD8α^+^ CD8β^+^ (of CD3^+^) | 11.7 ± 1.3 | 12 ± 1.1 | 13.2 ± 1.5 | 13.1 ± 1.2 | 13.8 ± 3.6 |
| NK cells (CD8α^+^CD3^-^ of lymphocytes) | 11.3 ± 0.9 | 11.4 ± 1.1 | 7.6 ± 1.3 | 10.1 ± 0.8 | 13.7 ± 1.7 |
| Macrophages (CD14^+^CD11b^-^CD163^+^ of leukocytes) | 2.3 ± 0.4 | 3 ± 1 | 3.3 ± 0.4 | 2.6 ± 0.9 | 3.6 ± 1 |
| Monocytes (CD14^+^CD11b^-^CD163^-^ of leukocytes) | 27 ± 2.6 | 27.6 ± 2.4 | 34.2 ± 4 | 36.9 ± 3.1 | 48.5 ± 9.3 |
| Neutrophils (CD14^+^CD16^+^CD163^-^ of leukocytes) | 10.5 ± 1.2 | 7.9 ± 0.9 | 12 ± 1.4 | 8.4 ± 1 | 9.4 ± 1.2 |

Table S6. Frequency (mean ± SEM) of leukocyte populations in blood at 20 days post vaccination

| Immune cell population | Group 1: Vaccinated *CD1D−/−* | Group 2: Vaccinated *CD1D−/+* | Group 3: Unvaccinated *CD1D−/−* | Group 4: Unvaccinated *CD1D−/+* | Group 5: Negative *CD1D−/+* ^a^ |
| --- | --- | --- | --- | --- | --- |
| CD3^+^ (of lymphocytes) | 56.3 ± 4.8 | 53.3 ± 4.1 | 63.5 ± 2.8 | 56.9 ± 1.9 | 68.9 ± 4.5 |
| αβ cells (CD3^+^TCRδ^-^ of lymphocytes) | 27.1 ± 2 | 31.1 ± 2.6 | 35.6 ± 2.2 | 34 ± 5.4 | 37.9 ± 3.8 |
| γδ cells (CD3^+^TCRδ^+^ of lymphocytes) | 26.4 ± 3.3 | 15.8 ± 2.4 | 21.1 ± 2.8 | 19.7 ± 3.4 | 18.7 ± 9.7 |
| CD4^-^CD8α^+^ (of CD3^+^) | 16.8 ± 1.4 | 17.3 ± 1.9 | 16.7 ± 1.6 | 18.6 ± 2.1 | 22.2 ± 4 |
| CD4^+^CD8α^+^ (of CD3^+^) | 12.6 ± 1.4 | 14 ± 1 | 13.1 ± 2.1 | 11.1 ± 1.3 | 12 ± 1.5 |
| CD4^+^CD8α^-^ (of CD3^+^) | 30.1 ± 2.1 | 37.4 ± 3.6 | 39.2 ± 1.4 | 38.5 ± 6.4 | 38.6 ± 7 |
| CD8α^+^ CD8β^+^ (of CD3^+^) | 14.5 ± 1.3 | 14.4 ± 1.9 | 14.2 ± 1.4 | 15.3 ± 1.8 | 18.6 ± 4.3 |
| NK cells (CD8α^+^CD3^-^ of lymphocytes) | 6.5 ± 1.9 | 8.1 ± 2 | 5.7 ± 0.8 | 8 ± 1.8 | 2.7 ± 0.4 |
| Macrophages (CD14^+^CD11b^-^CD163^+^ of leukocytes) | 2.2 ± 0.2 | 2.9 ± 0.6 | 1.7 ± 0.3 | 1.8 ± 0.2 | 3 ± 0.6 |
| Monocytes (CD14^+^CD11b^-^CD163^-^ of leukocytes) | 27.1 ± 3 | 29.2 ± 6.5 | 34.5 ± 3.8 | 39.3 ± 4.4 | 47.7 ± 0.5 |
| Neutrophils (CD14^+^CD16^+^CD163^-^ of leukocytes) | 9.2 ± 1.3 | 9.6 ± 1.1 | 12.5 ± 1.5 | 11.1 ± 2.1 | 9.2 ± 1.8 |

^a^ The blood of negative *CD1D*-/+ pigs were collected at 17 days post vaccination.

Table S7. Frequency (mean ± SEM) of leukocyte populations in blood at 5 days post challenge

| Immune cell population | Group 1: Vaccinated *CD1D−/−* | Group 2: Vaccinated *CD1D−/+* | Group 3: Unvaccinated *CD1D−/−* | Group 4: Unvaccinated *CD1D−/+* | Group 5: Negative *CD1D−/+* ^a^ |
| --- | --- | --- | --- | --- | --- |
| CD3^+^ (of lymphocytes) | 70 ± 5.4 | 69.2 ± 3.7 | 71.8 ± 4.9 | 71.7 ± 1.2 | 68.9 ± 4.5 |
| αβ cells (CD3^+^TCRδ^-^ of lymphocytes) | 42.7 ± 1.7 | 51.5 ± 4 | 36.7 ± 2.8 | 39.1 ± 3.8 | 37.9 ± 3.8 |
| γδ cells (CD3^+^TCRδ^+^ of lymphocytes) | 24.2 ± 3.2 | 12.2 ± 3.7 | 31.7 ± 3.5 | 31.3 ± 3.9 | 18.7 ± 9.7 |
| CD4^-^CD8α^+^ (of CD3^+^) | 20.9 ± 1.1 | 24.2 ± 2.9 | 17.4 ± 2.2 | 18.2 ± 1.3 | 22.2 ± 4 |
| CD4^+^CD8α^+^ (of CD3^+^) | 12.7 ± 1.7 | 17.3 ± 2.4 | 9.1 ± 1.3 | 8.5 ± 1.1 | 12 ± 1.5 |
| CD4^+^CD8α^-^ (of CD3^+^) | 34.5 ± 2.4 | 43.4 ± 5.9 | 28.2 ± 3.2 | 31.5 ± 4.8 | 38.6 ± 7 |
| CD8α^+^ CD8β^+^ (of CD3^+^) | 17.7 ± 1.1 | 20.4 ± 2.3 | 15.8 ± 2.8 | 14.9 ± 1.6 | 18.6 ± 4.3 |
| NK cells (CD8α^+^CD3^-^ of lymphocytes) | 2.6 ± 0.6 | 2.8 ± 0.6 | 2.4 ± 1.2 | 2.2 ± 0.3 | 2.7 ± 0.4 |
| Macrophages (CD14^+^CD11b^-^CD163^+^ of leukocytes) | 2.8 ± 0.3 | 4.3 ± 0.9 | 5.8 ± 1.6 | 5.5 ± 1.3 | 3 ± 0.6 |
| Monocytes (CD14^+^CD11b^-^CD163^-^ of leukocytes) | 33.7 ± 3.2 | 46.8 ± 4.5 | 28.1 ± 3 | 36.3 ± 3.1 | 47.7 ± 0.5 |
| Neutrophils (CD14^+^CD16^+^CD163^-^ of leukocytes) | 12.3 ± 2.5 | 13.7 ± 1.5 | 6.5 ± 0.8 | 10.8 ± 1.9 | 9.2 ± 1.8 |

^a^ The blood of negative *CD1D*-/+ pigs were collected at 17 days post vaccination.

Table S8. Shared expanded CDR3β clones between this study and prior study

| CDR3β | Number of clones in this study | Number of clones in Ref. (40) |
| --- | --- | --- |
| CASSGTGDIQYF | 2 | 3 |
| CASSNSYNSPLHF | 3 | 2 |
| CASSPGQYDYNF | 3 | 2 |
| CASSRDNSPLHF | 11 | 4 |
| CASSRDRDTDPLYF | 2 | 3 |
| CASSRDRGDTYFF | 2 | 5 |
| CASSRDRGTEVFF | 2 | 3 |
| CASSRDSYDYNF | 5 | 2 |
| CASSSAGQGTEVFF | 6 | 3 |
| CASSSELSQTQYF | 14 | 2 |
| CASSTGGYDYNF | 40 | 3 |
| CASSVGSYNDLHF | 6 | 2 |
| CGARGANTGQLYF | 6 | 2 |
| CGASDEDSYDYNF | 4 | 2 |
| CGASDRASRAAQLYF | 4 | 2 |
